# “Genetic and chemical disruption of Amyloid Precursor Protein processing impairs zebrafish sleep maintenance”

**DOI:** 10.1101/2022.06.08.495312

**Authors:** Güliz Gürel Özcan, Sumi Lim, Thomas Canning, Lavitasha Tirathdas, Joshua Donnelly, Tanushree Kundu, Jason Rihel

## Abstract

Amyloid precursor protein (APP) is a brain-rich, single pass transmembrane protein that is proteolytically processed into multiple products, including Amyloid-beta (Aβ), a major driver of Alzheimer’s disease (AD). Although both over-expression of APP, as in mouse models used to study AD, and exogenously delivered Aβ lead to changes in sleep behaviors, whether APP processing plays an endogenous role in regulating sleep is unknown. Here, we demonstrate that APP processing into Aβ40 and Aβ42 is conserved in zebrafish and then describe sleep/wake phenotypes in loss of function *appa* and *appb* mutants. Larvae with mutations in *appa* had reduced waking activity but normal sleep patterns, while larvae that lacked *appb* had shortened sleep bout durations at night. Treatment with the γ-secretase inhibitor DAPT also shortened sleep bouts at night, while the BACE-1 inhibitor lanabecestat lengthened sleep bouts. Since both drugs fail to further alter sleep in *appb* mutants, and intraventricular injection of the App cleavage product P3 also shortens night-time sleep bouts, we conclude that the proper balance of Appb proteolytic processing is required for normal sleep maintenance in zebrafish.

## Introduction

Sleep disturbances are prevalent in neuropsychiatric and neurodegenerative disorders such as Alzheimer’s Disease (AD). In addition to cognitive impairment, individuals with AD experience altered sleep patterns, including reduced rapid eye movement (REM) and non-REM sleep, increased wakefulness, sleep fragmentation, and electroencephalogram (EEG) abnormalities^1^. These sleep symptoms can manifest years before cognitive decline, and alterations in sleep can influence the deposition rates of Aymloid-β (Aβ) plaques that are the hallmark of AD, contributing to disease risk^2–4^. Consistent with this, some transgenic AD mouse models that overproduce Aβ display sleep disruptions prior to plaque formation, even without evident neuronal loss^5–11^. Recently, human studies have found that harbouring a high genetic risk for AD correlates with sleep changes, such as increased sleep rebound following sleep loss, even in young adults^12^. These observations suggest that there may be underlying early biological processes important for sleep regulation that are governed by AD susceptibility genes and contribute to disease progression.

One of the risk genes for AD encodes the Amyloid Precursor Protein (APP), a transmembrane protein that is proteolytically processed into multiple smaller fragments, including AD-associated fragments such as Aβ40 and Aβ42. Mutations in *App* are associated with both early and late onset AD^13,14^ including some alleles that are dominant and fully penetrant for early-onset AD^15^. Duplications of the App gene, including those associated with trisomy 21 (Down’s Syndrome), also confer increased AD risk^16^. However, APP is associated with other phenotypes beyond AD and has been ascribed a variety of other physiological roles in cell adhesion, axon growth, synapse formation and function, and intracellular signal transduction^17–19^. Additionally, APP is processed into fragments other than Aβ, including P3, APPsα, APPsβ, and the APP intracellular domain (AICD), but the endogenous roles of these peptides is poorly understood. Whether APP and its proteolytic fragments play a role in sleep regulation is unknown.

To explore the role of APP and its derivatives in sleep regulation, we introduced loss-of-function mutations into both zebrafish orthologs of the human APP gene, *appa* and *appb*, and evaluated their sleep-wake behaviours using a high-throughput behavioural assay^20^. Zebrafish are a good model to address this question because they possess the complete App processing machinery, including α-secretase (ADAM10), β-secretase proteins (BACE-1 and 2), and the components necessary for γ-secretase complex assembly, including Presenilin-1 (PSEN-1), Presenilin-2 (PSEN-2), Presenilin enhancer 2 (PEN-2), APH1 (anterior pharynx-defective 1), and Nicastrin (NCSTN)^21–25^. In addition, our previous work had revealed that depending on its structural configuration, Aβ can either increase or decrease sleep duration in larval fish by signalling through distinct receptors^26^, suggesting that App-derived products may act as sleep signals in zebrafish. By combining genetic analysis of *app* mutants with pharmacological interventions that block γ-secretase- and BACE-1-dependent cleavage of APP, we provide evidence that proteolytic processing of *appb* is required for maintaining sleep at night in zebrafish larvae.

## RESULTS

### Characterizing zebrafish *appa* and *appb* and generating mutants

There are two *app* genes in zebrafish, *appa* and *appb*, which raises the question whether they have redundant functions. We first examined their relationship to the human APP isoforms, because gene duplications often take on isoform or tissue-specific roles^27^. Zebrafish Appa is more similar to the Kunitz type protease inhibitor (KPI) domain-containing APP751 and 770 isoforms, while the Appb protein lacks the KPI domain and is more similar to the APP695 isoform (Figure 1A and B)^28,29^. Both Appa and Appb have respectively an 80% and 71% conserved identity in the Aβ42 region compared to human APP, with conserved proteolytic cleavage sites for processing by α, β, and γ secretases (Figure 1B). Both *appa and appb* genes are abundantly expressed in the zebrafish brain, although with non-overlapping expression patterns (Figure1C). For example, while *appa* is expressed strongly in the cerebellum caudal lobe, olfactory bulb and torus longitudinalis, *appb* is more strongly expressed in nuclei in the hindbrain (Figure1C, Figure S1 and S2). Examination of the brain expression pattern and levels of *appa* and *appb* during the day (4 hours post lights on) and night (4 hours post lights off) failed to detect any time-of-day differences, either globally or region specifically (Figure S2B and Figure S3D). *appb* is also highly expressed in the very early stages of zebrafish development, indicating that it is maternally deposited (Figure S3A) Together, the gene expression patterns and structural homology differences of zebrafish Appa and Appb are consistent with these proteins possibly having both isoform and brain tissue-specific functions.

**Figure 1.**
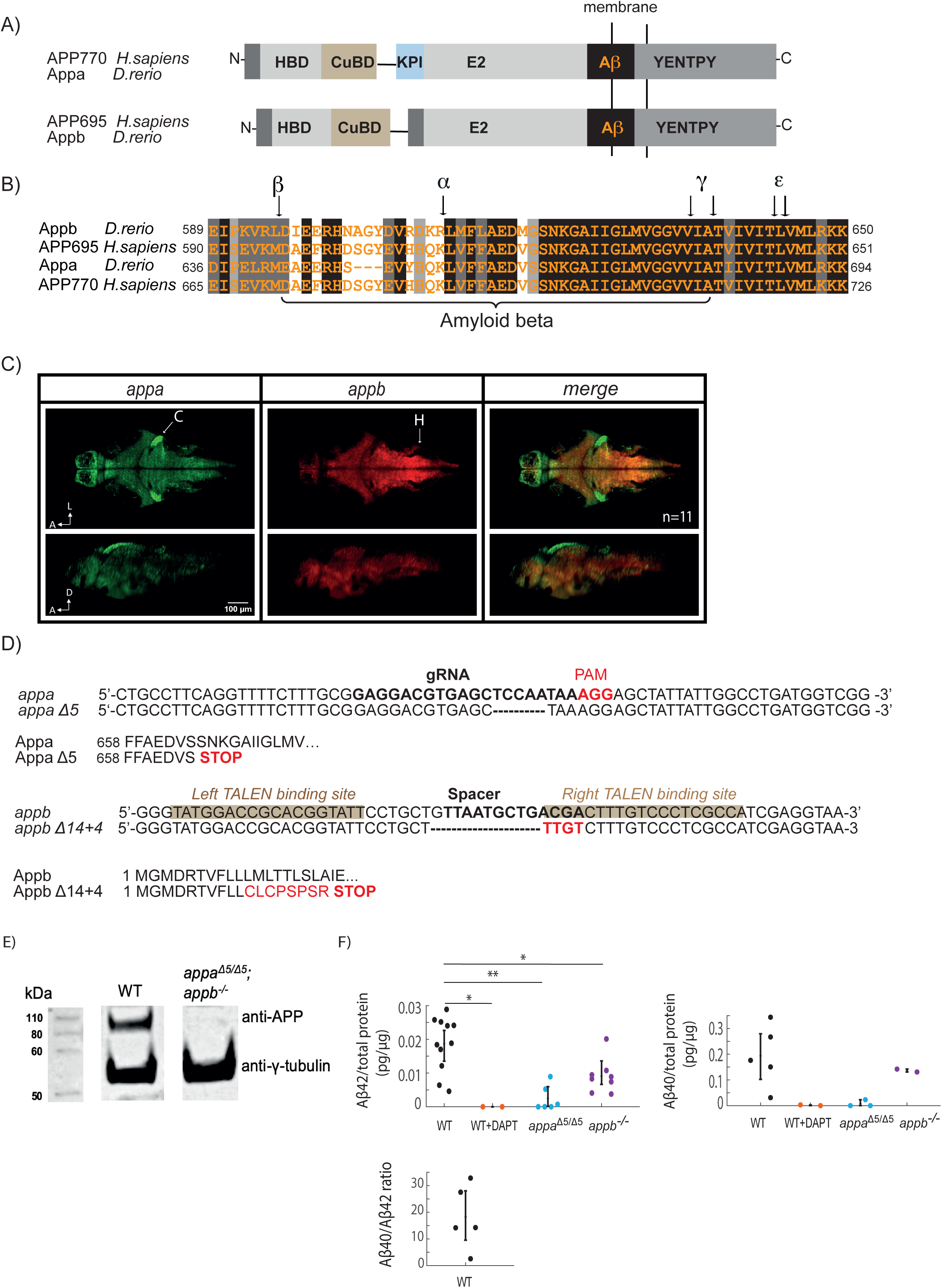
Zebrafish App protein organisation, gene expression, and mutant generation. **A)** There are 2 *App* orthologs, *appa* and *appb* in zebrafish. Appa contains the Kunitz type protease inhibitor (KPI) domain and thus has a similar gene organization to the human APP770 isoform. Zebrafish Appb lacks the KPI domain similar to the brain enriched human APP695 isoform. Both Appa and Appb have the functional App domains, including the Heparin binding domain (HBD), Copper binding domain (CuBD), extracellular E2 domain (E2), the conserved YENTPY motif, and amyloid beta region (AB) **B)** Alignment of Aβ regions of zebrafish Appa and Appb to human APP695 and APP770 shows high conservation within the Aβ region and the proteolytic cleavage sites (indicated with black arrows): α-secretase cleavage site (α), β-secretase cleavage site (β), γ-secretase cleavage sites (γ) and ε-cleavage sites (ε). Black, dark grey, and light grey boxes indicate strictly, highly, and moderately conserved amino acid residues, respectively. **C)** As detected by multiplexed hybridization chain reaction (HCR), *appa* (green) and *appb* (red) are both expressed widely but in non-overlapping regions of the 5dpf larval brain, including the cerebellum and nuclei in the hindbrain (C=cerebellum, H=hindbrain). Shown is a representative image from a single brain taken at one z-plane (z140/420) (dorsal view, above) and through the midline of the same brain (lateral view, below) from an experiment with total of n=22 fish. A=Anterior, D=dorsal, L = Left. **D)** CRISPR/Cas9 targeting of zebrafish *appa* resulted in a 5 bp deletion. The target gRNA sequence is shown in bold, and the obligatory PAM sequence (AGG) is in red. The predicted translation of *appaΔ5* leads to a premature stop codon within the Aβ region (Appa amino acid 665). TALEN targeting of zebrafish *appb* resulted in a 14 bp deletion (dashes) and 4 bp insertion (red). The left and right sites targeted by the TALENs are highlighted, and the spacer sequence, where cleavage occurs, is bolded. The predicted translation of *appbΔ14+4* leads to a frameshift and a premature stop codon. **E)** Western blot analysis of APP in brain homogenates from wildtype (WT) and *appa^Δ5/Δ5^; appb^-/-^* double mutants **F)** Elisa detection of Aβ42 (left) and Aβ40 (right) levels in adult brain homogenates from WT controls, WT animals treated with the γ-secretase inhibitor DAPT for 24 hrs, *appa^Δ5/Δ5^*, and *appb^-/-^* mutants quantified as pg of Aβ per µg total protein extracted. Each dot is an independent biological replicate. The bottom panel plots the ratio of Aβ40 to Aβ42 in WT zebrafish adult brain homogenates, n=5. *p≤0.05, **p≤0.01, Dunnett’s test.

We next generated zebrafish mutants with deletions in the *appa* and *appb* genes. To isolate mutations in *appa*, we used CRISPR/Cas9^30^.The CRISPR design tool CHOPCHOP (http://chopchop.cbu.uib.no) was used to identify candidate gRNAs to target the conserved Αβ region (amino acid residues 25-35) of *appa*^31^, which was then co-injected with Cas9 mRNA into zebrafish eggs at the one cell-stage. Injected animals (F0s) that harbored frameshift mutations were then identified by Illumina sequencing and outcrossed to wild-type animals to generate mutant families (see Methods). One family was isolated that harbors a 5 base pair frameshift deletion (*appa^Δ^*^5^) that leads to an early stop codon within the Aβ domain. The *appa^Δ^*^5^ allele therefore lacks the conserved residues 26-42 of the Aβ and the entire intracellular C-terminus of Appa (Figure 1B, D).

To generate a loss of function mutation in the *appb* gene, two Transcription activator-like effector nuclease (TALEN) arms targeting a conserved region within the first exon of the zebrafish *appb* gene were designed using the Zifit software (http://zifit.partners.org/ZiFiT/)^32^.These TALEN arms were each fused to one half of a Fok1 heterodimer to generate mutagenic double strand breaks within the first exon of *appb* (Figure 1D). F0 fish that had been co-injected at the one cell stage with mRNA encoding the two TALEN arms were Illumina sequenced to identify a founder that contains an *appb* allele (*appb^Δ14+^*^4^) with a 14 base pair deletion and a 4 base pair insertion that generates a frameshift followed by an early stop codon. This founder was used to generate a stable heterozygous family (herein called *appb*^-/+^) for subsequent behavioral analysis (Figure 1D).

To confirm that these *appa* and *appb* alleles represent loss of function mutations, we performed Western blot analysis on brain homogenates from adult double homozygous *appa ^Δ^*^5^*^/Δ5^*; *appb*^-/-^ mutants using an anti-APP antibody (22C11) that recognizes both zebrafish Appa and Appb. APP protein was detectable as a ∼100 kD band in WT but not in *appa^Δ5/Δ5^*; *appb*^-/-^ double mutants, confirming that neither Appa nor Appb are made (Figure 1E and Figure S3C). We also observed by qRT-PCR that mutant *appb* transcripts are only 25% of WT levels, consistent with non-sense mediated decay that is often observed in transcripts harboring an early termination codon (Figure S3B)^33^.

Although zebrafish have the machinery to process App into Aβ fragments, whether Aβ40/42 are actually generated in zebrafish has not been formally demonstrated. To measure endogenous Aβ levels, we used a highly sensitive electrochemiluminescence based ELISA kit and detected in adult brains ∼0.02 pg Aβ42/μg total protein and ∼0.2 pg Aβ40/μg total protein, yielding an Aβ42:Aβ40 ratio of 1:15-1:20 (Figure 1F), similar to the range observed in various mammalian species^34^. Aβ40 and Aβ42 were completely undetectable in brains following overnight exposure of WT adults to the y-secretase inhibitor DAPT, confirming the efficacy of this drug to block Aβ40/42 production in zebrafish (Figure 1F). Moreover, levels of both Aβ40 and Aβ42 in either *appa^Δ5/Δ5^* or *appb^-/-^* mutants were significantly lower than in WT animals, suggesting that Aβ is made from both Appa and Appb (Figure 1F). The reduction of Aβ levels in *appa^Δ5/Δ5^* mutants (−89% and −90% for Aβ40 and Aβ42, respectively) was stronger than in *appb^-/-^* mutants, which may suggest that more Aβ is generated from Appa than from Appb; however, the Aβ40/42 epitopes detected by this kit are better matched in the Appa sequence than Appb, so the relative difference in detected Aβ levels may be due to slight differences in antibody affinity/detection between Appa and Appb. Nevertheless, these results are consistent with Aβ being produced from both Appa and Appb in zebrafish, and demonstrate that both mutants have disruptions in Aβ production.

### appa^Δ^^5/Δ^^5^ and appb^Δ14+4^ (appb^-/-^) have distinct sleep-wake profiles

Zebrafish *appa^Δ5/Δ5^* mutants do not have any obvious morphological abnormalities during development, have normal survival rates to adulthood, and are generally healthy and fertile. To examine whether *appa^Δ5/Δ5^* mutant larvae have sleep or wake phenotypes, we used automated video monitoring to track larvae from in-crosses of *appa^Δ^*^5^*^/+^* parents over several days on a 14hr:10hr light:dark cycle (Figure 2). Sleep in zebrafish is defined as a period of inactivity lasting longer than one minute, as quiescent periods lasting at least this long are associated with an increased arousal threshold and other features of behavioral sleep, including circadian and homeostatic regulation^35^. Zebrafish sleep is organized into bouts, with the sleep bout length describing the duration of consecutive, uninterrupted minutes of sleep. We also measured the vigor of their movements during the active bouts, quantified as the average waking activity. Sleep and waking activity are therefore not identical and can be selectively and differentially modulated by drugs^20^ or mutation^36^. Assessing these parameters across the day and night for *appa^Δ5/Δ5^* mutants and their wild-type (WT) siblings uncovered subtle differences in mutant behavior. The *appa^Δ5/Δ5^* mutant had a reduction of 9.0% (lower bound, −15.0%; upper bound, −3%, 95% confidence interval, CI) in waking activity during the day compared to *appa^+/+^* siblings (Figure 2A and 2B). At night, *appa^Δ5/Δ5^* mutants also had slightly lower waking activity levels (−4.7%, [-10.0; −0.5, 95%CI]) (Figure 2C). In contrast, neither the total sleep (Figure 2E and 2F) nor the structure of sleep, such as the number and duration of sleep bouts (Figure S4) were statistically different across genotypes during either the night or the day.

**Figure 2.**
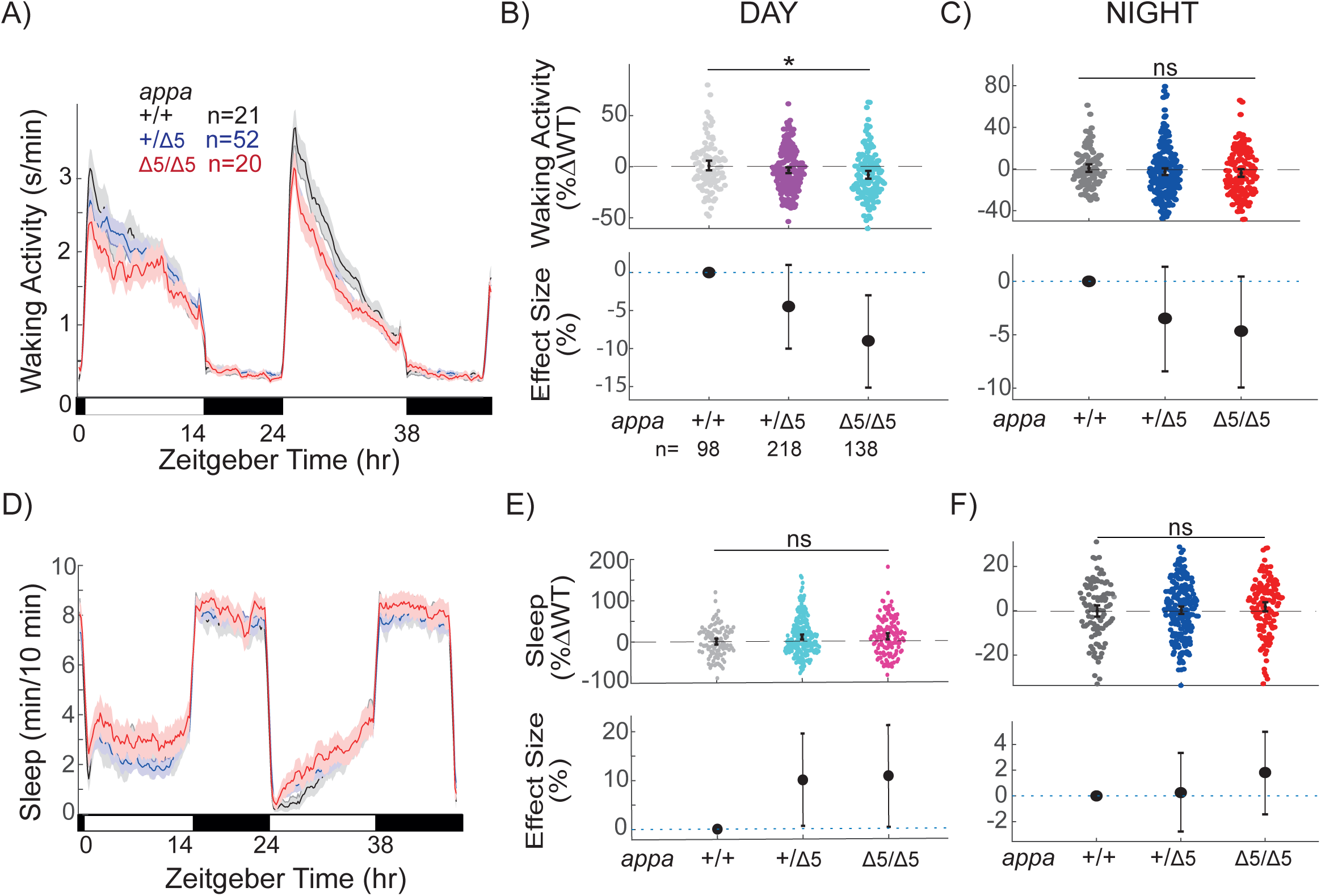
*appa^Δ5/Δ5^* mutants have reduced day waking activity but no sleep phenotype. Exemplar 48 hr traces of average waking activity taken from a single experiment of *appa^Δ5/Δ5^* mutants, heterozygous, and wild type siblings (5-7dpf) on a 14hr:10hr light:dark cycle. Each line and shaded ribbon show the mean ± SEM. **B**) Day waking activity and **C)** night waking activity for *appa^Δ5/Δ5^* mutants and siblings from *appa^+/Δ^*^5^ incrosses, combined across N=5 experiments. Each dot is a single larva, normalized to the mean of their experimentally-matched WTs. At the bottom are plotted the effect sizes (± 95% confidence interval, CI) relative to WT. **D)** Exemplar 48 hr traces of average sleep for the same experiment shown in A. **E)** Day sleep and **F**) night sleep of WT, heterozygous, and *appa^Δ^*^5^*^/ Δ^*^5^ mutants normalized to their WT siblings, as in B and C. ^ns^p>0.05, *p≤0.05, Kruskal-Wallis, Tukey’s post hoc test. n= number of larvae.

Together, these data show Appa is not required for normal sleep states in zebrafish larvae, although it influences locomotor drive during the waking day.

We also did not observe any obvious developmental delays or morphological abnormalities in *appb^-/-^* mutant larvae or adults and therefore assessed whether Appb might play a non-redundant role in larval sleep regulation. Similar to *appa^Δ5/Δ5^* mutants, *appb^-/-^* larvae had a reduction in day-time waking activity of 13.0% [-20.2; − 5.6, 95%CI] relative to *appb*^+/+^ siblings (Figure 3A, B). However, unlike *appa^Δ5/Δ5^* animals, *appb^-/-^* larvae had an increase in activity of 8.2% [3.3; 13.0, 95%CI] specifically at night (Figure 3C). We also observed that while *appb^-/-^* larvae had unaffected sleep during the day (Figure 3D, E), they had a 7.9% (−14.0; −7.0, 95%CI) reduction in sleep at night (Figure 3F), which corresponds to ∼30 min less sleep per night. Thus, both Appa and Appb regulate daytime waking activity levels but have non-overlapping roles in regulating nighttime activity and sleep, with only *appb* mutants exhibiting sleep phenotypes.

**Figure 3.**
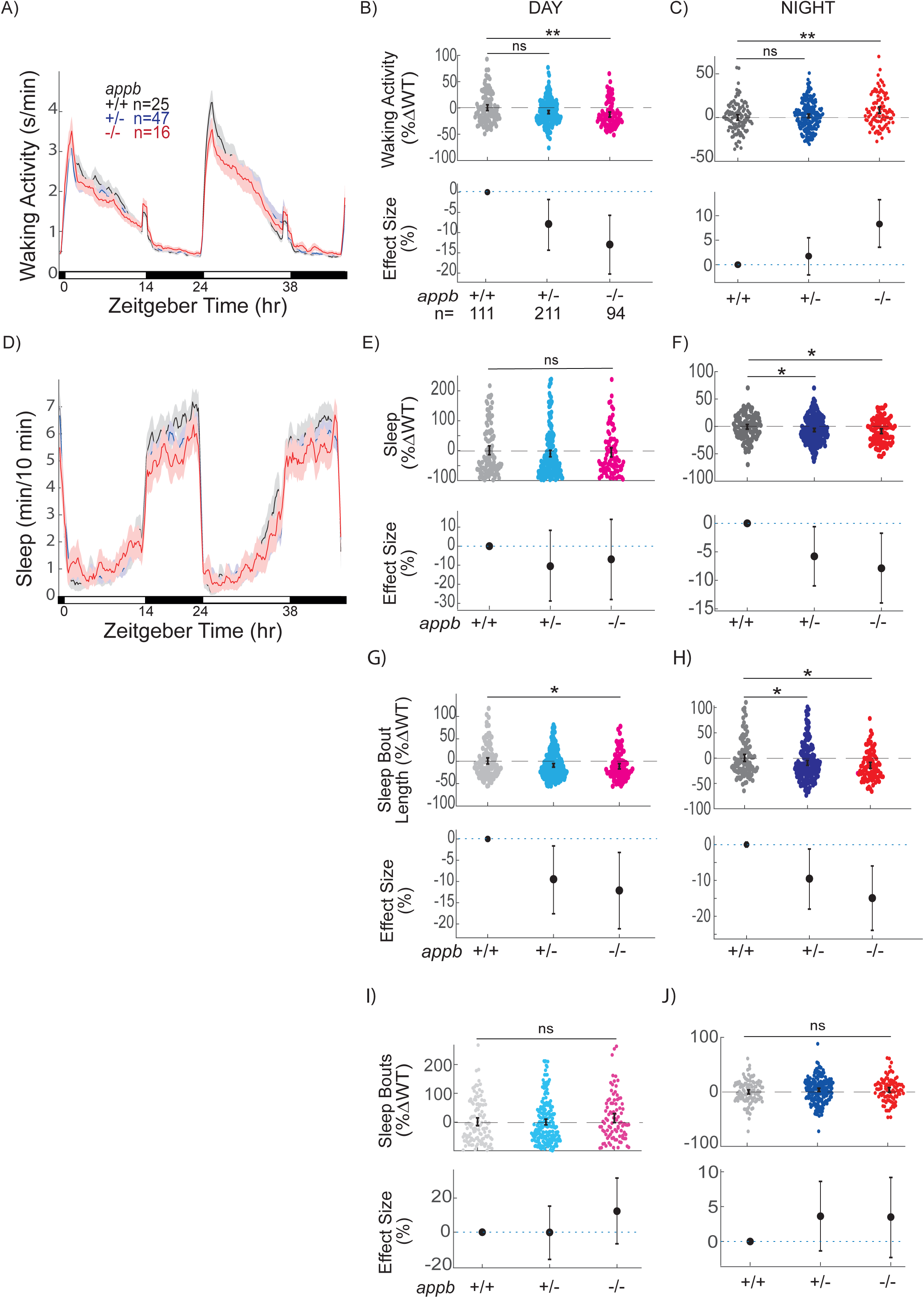
*appb^-/-^* mutants have altered waking activity and sleep across the day-night cycle. Exemplar 48 hr traces of average waking activity taken from a single experiment of *appa^Δ^*^5^ mutants, heterozygous, and wild type siblings (5-7dpf) on a 14hr:10hr light:dark cycle. Each line and shaded ribbon show the mean ± SEM. **B**) Day waking activity and **C)** night waking activity for *appb^-/-^* mutants and siblings from *appb^+/-^* incrosses, combined across N=5 experiments. **D)** Exemplar 48 hr traces of average sleep for the same experiment shown in A. **E)** Day sleep, **F**) night sleep, **G)** day sleep length, **H)** night sleep length, **I)** day sleep bout number, and **J)** night sleep bout number of WT, heterozygous and *appb^-/-^* mutants normalized to WT siblings as in B and C. At top, each dot is an animal normalized to the mean of their experimentally-matched WT. At bottom, shown are the effect size ± 95%CI relative to WT. ^ns^p>0.05, *p≤0.05, **p≤0.01, Kruskal-Wallis, Tukey’s post hoc test. n= number of larvae.

To further investigate the nature of the decreased night sleep in *appb^-/-^* mutants, we compared the sleep architecture of these mutants to their wild-type and heterozygous siblings. We specifically examined whether the change in total sleep was due to alterations in the number of sleep bouts (i.e., how often sleep is initiated) or in the average lengths of sleep bouts (i.e., once sleep is initiated, how long it is maintained). The average sleep bout length was shorter by 12.1% (−21.2; −3.2, 95%CI) in *appb^-/-^* mutants during the day and by 14.9% (−23.9; −5.9, 95%CI) at night compared to their WT siblings (Figure 3G, H). The number of sleep bouts during either the day or night were not significantly different between *appb^-/-^* mutants and wild type animals (Figure 3I, J). These results show that the *appb^-/-^* mutants initiate sleep normally but cannot sustain continuous sleep as long as WT, indicating a defect in sleep maintenance.

### γ- and β-secretase inhibitors modulate sleep maintenance in an Appb dependent manner

Because App undergoes complex proteolytic processing, we decided to test whether drugs that inhibit App cleavage also modulate sleep in an Appb dependent manner. We first tested whether the γ-secretase inhibitor DAPT, which effectively prevented Aβ production in zebrafish after 24 hrs (Figure 1F), alters sleep and waking activity in ether WT or *appb* homozygous mutants (Figure 4A-D). Unlike either *appa* or *appb* mutants, which had lower waking activity during the day, DAPT significantly increased WT waking activity, with no effect at night (Figure 4A, Figure S5A,E). In contrast, DAPT significantly reduced daytime waking activity in *appb*^-/-^ larvae (Figure 4B, Figure S5A). Together, these results indicate that γ-secretase dependent cleavage products of Appb increase daytime waking, while other γ-secretase targets have a net effect of reducing wake activity.

**Figure 4.**
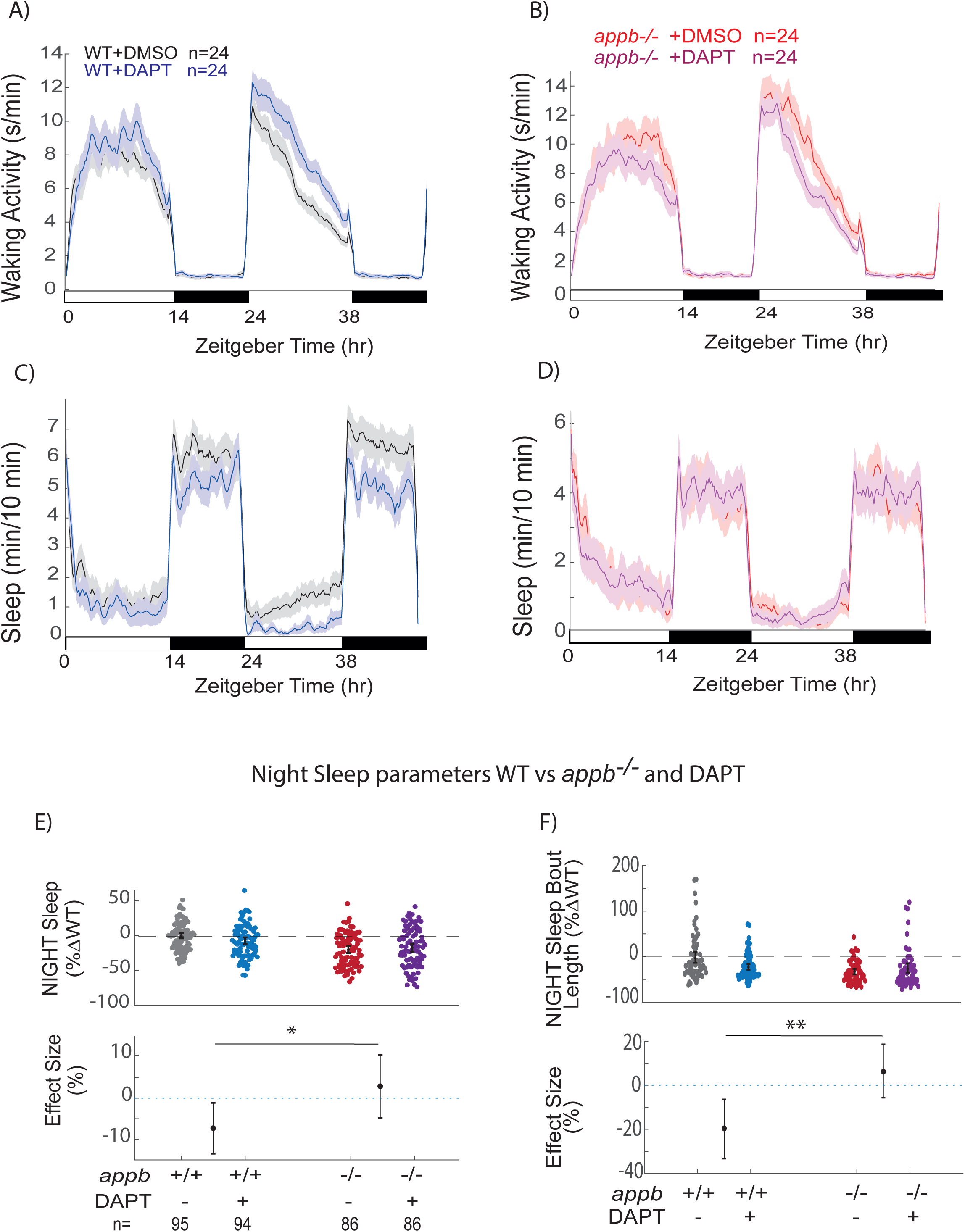
The y-secretase inhibitor DAPT shortens sleep bout lengths at night in WT but not in *appb*^-/-^ mutants. **A,B)** Exemplar 48 hr traces on a 14hr:10hr light:dark cycle of the average waking activity of WT (A) and *appb*^-/-^ mutants (B) continuously exposed to either 100 µM DAPT or DMSO vehicle control. **C,D)** Exemplar 48 hr traces of the average sleep from the same experiment shown in A and B. **E)** Night sleep and **F)** Night sleep bout length of WTs and *appb*^-/-^ mutants exposed to either 100 µM DAPT or DMSO vehicle. At top, each dot represents a single larva normalized to its experiment-matched WT (−DAPT) mean; error bars indicate ± SEM. At bottom, the within-genotype effect size and 95%CIs of DAPT treatment are plotted. n = the number of larvae. Data is pooled from N =4 independent experiments, omitting the first day and night to account for any delay in drug action. ^ns^p>0.05, *p≤0.05, **p≤0.01, 2-way ANOVA, Tukey’s post hoc test. n= number of larvae.

DAPT also significantly reduced total nighttime sleep of WT larvae by 7.29% (−13.38; −1.06, %95 CI) and trended to lowered daytime sleep by 18.46% (−38.58; +0.87, %95 CI) (Figure 4C, E and Figure S5B). This overall nighttime sleep reduction was due to a shortening in the average length of sleep bouts (−19.6% [-33.83; −6.55, %95CI], an effect size similar to *appb^-/-^* alone), even though the number of sleep bouts were slightly increased by DAPT (Figure 4F, Figure S5F). However, when DAPT was tested on *appb*^-/-^ mutants, there was no effect on total sleep or sleep bout lengths (Figure 4D-F). This significant, non-additive effect (genotype x drug interaction, p<0.05 for night sleep and p<0.01 for sleep bout length at night, two-way ANOVA) cannot be explained by a flooring effect, as wake-promoting drugs can reduce zebrafish larval sleep much more than observed in *appb^-/-^* mutants alone^20,37^. Instead, this demonstrates that DAPT requires the presence of Appb to influence sleep length at night, and further suggests that the short sleeping phenotype of *appb* mutants is due to the loss of Appb-derived γ-secretase cleavage products, such as Aβ, P3, or AICD.

To further dissect which Appb cleavage products modulate sleep and wakefulness, we next tested the effects of the BACE-1 inhibitor, lanabecestat, on WT and *appb*^-/-^ mutant behavior. Like γ-secretase inhibitors, blocking β-secretase will prevent Aβ production; however, other products blocked by γ-secretase inhibitors will remain unaffected, such as AICD, or even enhanced, such as P3, allowing us to test which Appb-derived products are responsible for shortening sleep (Table 1). Unlike DAPT but similar to *appa* and *appb* mutants, lanabecestat slightly reduced daytime waking activity of WT larvae (Figure 5A, Figure S6A), and similar to *appa* mutants, reduced daytime waking activity at night (Figure S6A,E). However, when *appb* mutants were exposed to lanabecestat, there was no longer an effect on daytime waking levels, while nighttime waking activity was even slightly increased (Figure 5B, Figure S6E). Thus, Appb must be present for β-secretase inhibition to exert an effect on daytime waking activity.

**Figure 5.**
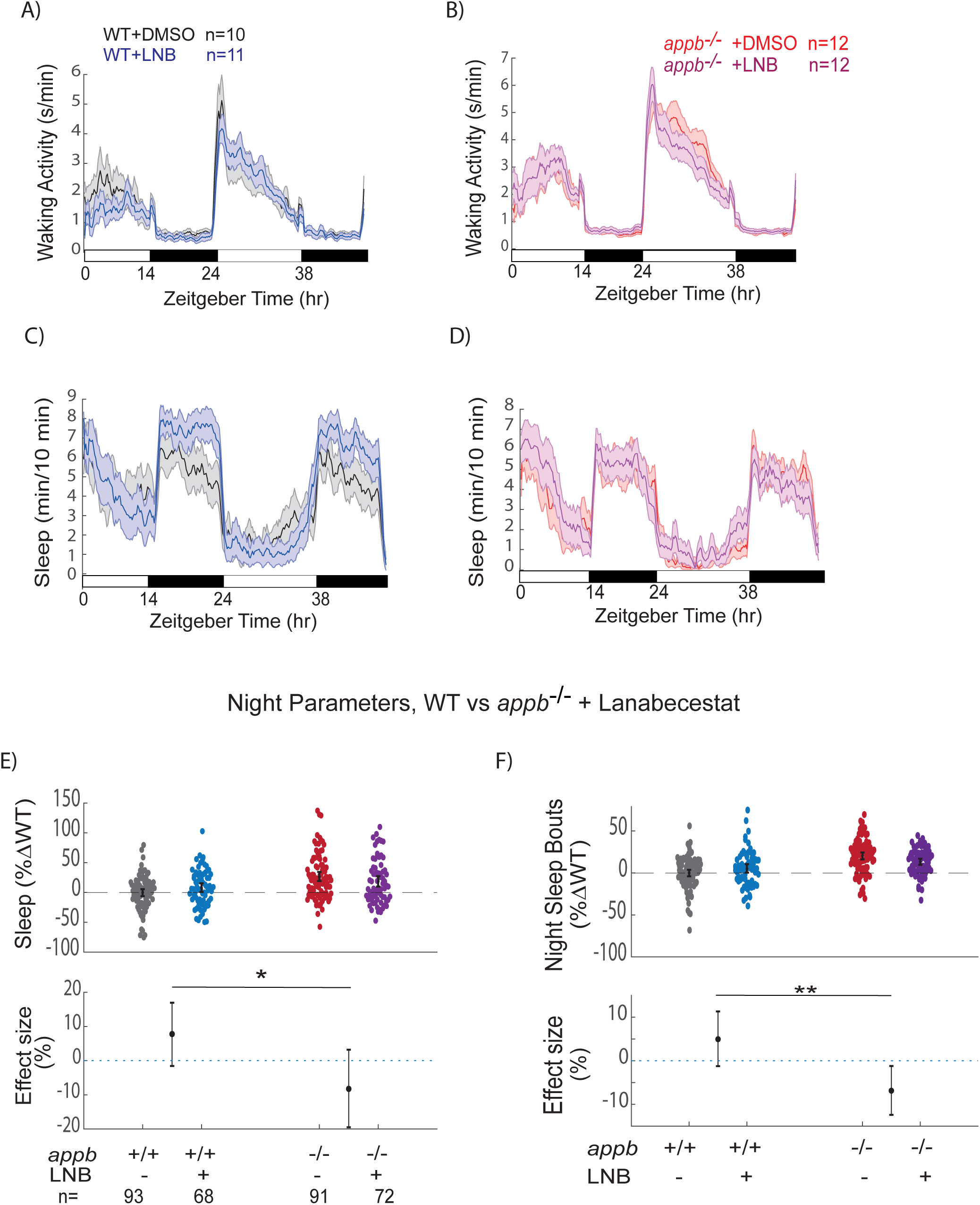
The β-secretase inhibitor Lanabecestat increases sleep at night in WT but not in *appb*^-/-^ mutants. **A,B)** Exemplar 48 hr traces on a 14hr:10hr light:dark cycle of the average waking activity of WT (A) and appb^-/-^ mutants (B) continuously exposed to either 0.3 µM Lanabecestat or DMSO vehicle control. **C, D)** Exemplar 48 hr traces of the average sleep from the same experiment shown in A and B. **E)** Night sleep and **F)** night sleep bout length of WTs and *appb*^-/-^ mutants exposed to either 0.3 µM Lanabecestat or DMSO vehicle. At top, each dot represents a single larva normalized to its experiment-matched WT (−Lanabecestat) mean, and error bars indicate ± SEM. At bottom, the within-genotype effect size and 95%CIs of 0.3 µM Lanabecestat treatment are plotted. n = the number of larvae. Data is pooled from N =4 independent experiments, omitting the first day and night to account for any delay in drug action. ^ns^p>0.05, *p≤0.05, **p≤0.01, 2-way ANOVA, Tukey’s post hoc test. n= number of larvae.

**Table 1.** Effects of blocking γ-secretase or β-secretase on proteolytic cleavage products on App and effects on night sleep. (↓, decrease; ↑, increase; X, blocked production; −, no effect)^38–40^.

| | APP | sAPP<br>$\alpha$ | C83 | P3 | AICD | sAPP<br>$\beta$ | C99 | Aβ | Night Sleep |
| --- | --- | --- | --- | --- | --- | --- | --- | --- | --- |
| *appb*<sup>-/-</sup> | x | x | x | x | x | x | x | x | ↓ |
| γ-secretase inhibition | ↑ | ↑ | ↑ | x | x | x | ↑ | x | ↓ |
| β-secretase inhibition | ↑ | ↑ | ↑ | ↑ | — | x | x | x | ↑ |

Furthermore, unlike DAPT and *appb* mutants, lanabecestat increased sleep at night in WT larvae (+7.78, [-1.37; +17.06, %95CI,]) (Figure 5C,E), due to an increase in both the number of sleep bouts (+4.95, [-6.34; +6.50, %95CI,]) and the average length of sleep bouts (+8.32, [-3.39; +20.45, %95CI]) (Figure 5E,F, and Figure S6F). Similarly, lanabecestat was unable to increase total sleep, the number of sleep bouts, or the sleep bout lengths of *appb*^-/-^ mutants at night (Figure 5D-F and Figure S6F). The significant non-additive interaction between lanabecestat and *appb* genotype (genotype x drug interaction, p<0.05 for night sleep and p<0.01 for sleep bout number at night, two-way ANOVA) suggests that inhibition of β-secretase requires the presence of Appb to influence sleep at night.

### P3 brain injections reduces the lengths of sleep bouts at night

Blocking either β-secretase or γ-secretase resulted in Appb-dependent, but opposing, sleep phenotypes at night. Since β- and γ-secretase inhibition differentially alter the formation of App cleavage products, of which all are lost in *appb*^-/-^ mutants, we hypothesized that the night-time sleep phenotypes might be explained by fragments that are blocked by γ-secretase inhibitors but enhanced or unchanged by β-secretase inhibition (Table 1). We therefore focused on P3, a partial Aβ fragment (Aβ_17-42_) that is boosted by BACE-1 inhibitors and absent when γ-secretase is blocked or *appb* is mutated. Injection of P3 into the brain ventricle of WT larvae had no effect on waking activity in either the day or the night (Figure 6A-C) but caused a significant 10.03% (−15.05; −5.17, %95CI) decrease in night sleep relative to vehicle-injected controls (Figure 6D & 6F). As in other manipulations of App processing, this nighttime reduction in sleep was caused by shortened sleep bout lengths (−16.50% [-26.60; −6.30, %95 CI]) rather than a change in the number of sleep bouts (Figure 6G-J). Although this demonstrates that App cleavage products such as P3 can have acute effects on sleep maintenance at night, since this effect is in the same direction as γ-secretase inhibition, which blocks the formation of P3, and in the opposite direction from BACE-1 inhibition, which enhances P3 production, alterations to P3 levels alone cannot explain the sleep phenotypes seen in *appb* mutants or drug manipulations that affect APP processing.

**Figure 6.**
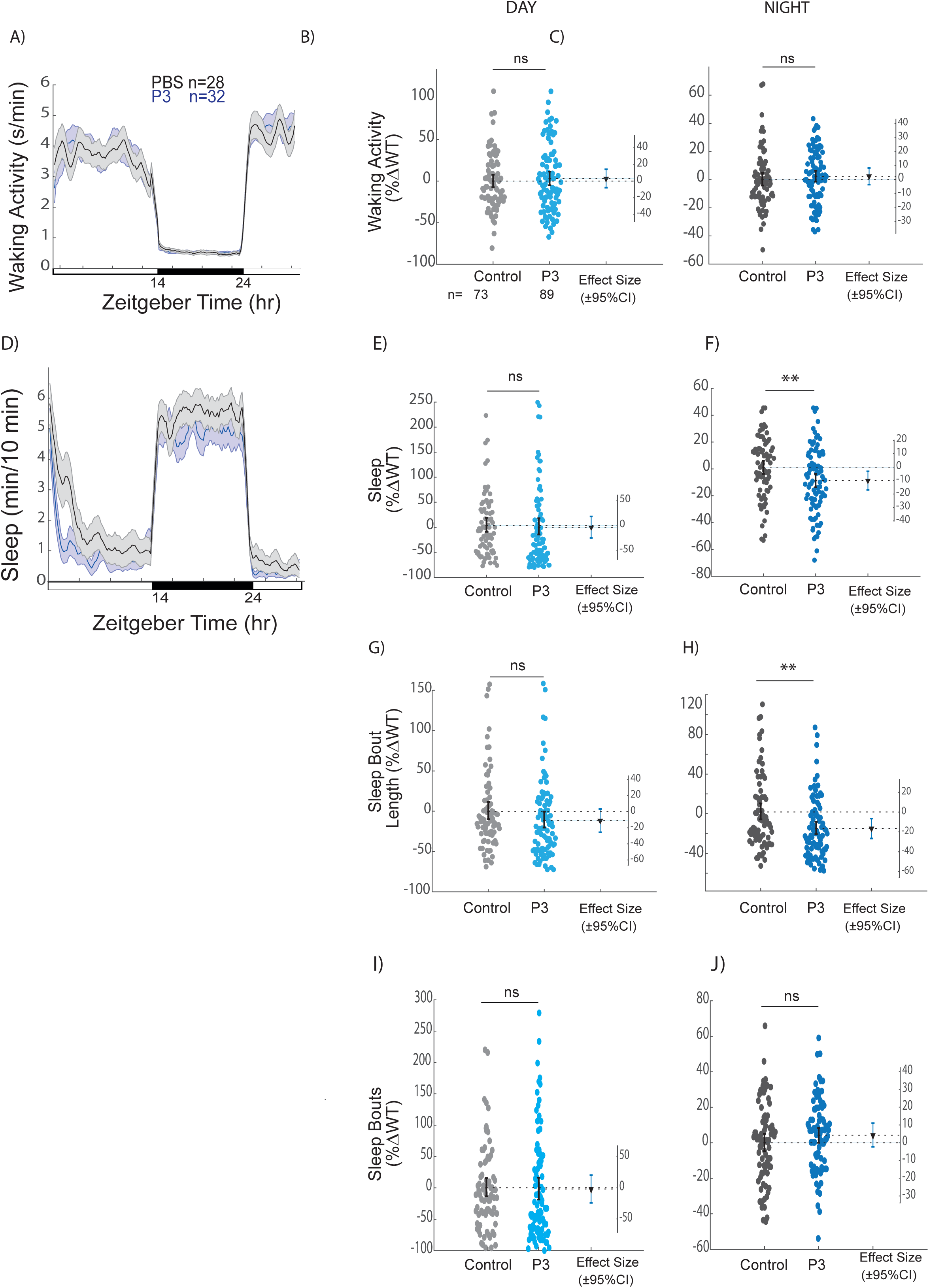
Intraventricular injection of P3 decreases sleep and shortens sleep bout lengths. Exemplar 24 hr traces of the average waking activity of WT larvae injected with either P3 or vehicle control on a 14hr:10hr light:dark cycle. Shown is a single exemplar experiment. **B,C)** Average waking activity during the day (B) and night (C) across 3 independent experiments. Each dot represents a single larva normalized to the experiment-matched WT mean. The bars represent the mean. **D)** Exemplar 24 hr trace of the average sleep for the experiment shown in A. **E)** Day sleep, **F)** night sleep, **G)** day sleep bout length, **H)** night sleep bout length, **I)** day sleep bout number, and **J)** night sleep bout number. n = the number of larvae. ^ns^p>0.05, *p≤0.05, **p≤0.01, Kruskal-Wallis, Tukey’s post hoc test. n= number of larvae.

## DISCUSSION

### Comparison to other App loss of function studies

We found that both *appa^Δ5/Δ5^* and *appb^-/-^* zebrafish had reduced locomotor activity during the day, but only *appb^-/-^* animals had a reduction in sleep maintenance at night. Previous studies of mouse *App* knockouts have also observed reduced locomotor and exploratory activity, but many other phenotypes have been described, including reduced growth and brain weight, reduced grip strength, hypersensitivity to seizures, alterations in copper and lipid homeostasis, reactive gliosis, and impaired spatial learning^41–50^. However, the lack of major morphological and developmental phenotypes in both *appa* and *appb* mutants stands in stark contrast to several reports that investigated zebrafish App function with morpholino knockdowns. For example, morpholino knockdown of zebrafish Appb has variously been reported to affect convergent-extension during gastrulation^51^, axon outgrowth of spinal motor neurons^52,53^, hindbrain neurogenesis^54^, or cerebrovascular development^55^. Our observations are in more in line with other recent studies that do not find large developmental phenotypes for zebrafish *app* mutants^56–58^. Since *appb*^-/-^ larvae from *appb*^-/-^ mothers also are morphologically normal (Figure 4-5), the difference in phenotype compared to morpholino studies cannot be explained by differential effects on maternally deposited transcripts of *appb* (Figure S3).

Although sleep has not been investigated in zebrafish *app* mutants before, two other studies have examined the role of *appb* on larval locomotor activity, coming to different conclusions than what we draw here. One *appb* morpholino study found knockdown resulted in hyperlocomotion between 28-45 hpf^52^, while we find both *appa^Δ5/Δ5^* and *appb*^-/-^ larvae at older stages (4-7dpf) are less active than their wild type siblings. These contrasting results are likely due to differences in locomotor behavior regulation at different stages of development, methodological differences (morpholino vs. knockout), or both. Another study of *appb*^-/-^ larvae tracked locomotor behavior for 60 minutes at a similar developmental age to our study (6 dpf) but found no differences in locomotion when comparing *appb* mutants to non-sibling wild type animals^56^. Given that we found *appa* and *appb* mutants are only 8-10% less active than sibling-matched wild type animals during the day, the short, 60 min observation window may not have been sufficient to capture this difference; alternatively, the time of day of observation might affect the ability to detect locomotor phenotypes (e.g., see Figure 2A and 3A). Indeed, we found *appb* mutants were significantly more, not less, active at night (Figure 3C).

Overall, our sleep and wake analysis of *appa* and *appb* mutants are broadly consistent with other rodent and zebrafish studies and expands the known phenotypes associated with App loss of function mutations. These results also demonstrate that Appa and Appb play at least partially non-redundant roles in the regulation of sleep and locomotor activity in larval zebrafish, which may explain why the zebrafish phenotypes of single mutants are somewhat milder than that reported for rodent App knockouts. Differences in the phenotypes between *appa* and *appb* mutants may reflect their differential expression patterns in the brain (Figure 1C and Figure S1 and S2A), overall expression levels, or possibly different sensitivities or exposure to App processing enzymes, yielding different ratios of cleavage products from Appa versus Appb. For example, we found that Appa may be a larger source of Aβ in larvae, as *appa^Δ5/Δ5^* mutants had lower detectable levels of Aβ compared with *appb*^-/-^ mutants (Figure 1F).

### Sleep maintenance and Appb proteolytic cleavage

We observed no sleep phenotypes in *appa^Δ5/Δ5^* mutants, but *appb^-/-^* mutants had reduced nighttime sleep due to an inability to maintain longer sleep bout durations. Inhibition of gamma-secretase also shortened the length of nighttime sleep bouts in an Appb-dependent manner, suggesting that gamma-secretase dependent cleavage products of Appb such as Aβ, P3, or AICD may act as a signal for maintaining nighttime sleep that is lost in *appb*^-/-^ mutants. Previous work has demonstrated that exogenously delivered Aβ does have both sleep-promoting and inhibiting properties when injected into zebrafish larvae^26^, engaging many areas of the brain including the sleep-promoting galanin-positive neurons of the preoptic area and hypothalamus^59^. However, that work found that longer Aβ oligomers promoted sleep predominantly by increasing sleep initiation rather than altering sleep bout maintenance, while shorter forms of Aβ promoted wakefulness^26^. Moreover, inhibition of BACE-1, which also prevents the formation of Aβ, instead led to increased sleep maintenance in an Appb dependent manner. This suggests that loss of Appb-cleavage products other than Aβ are responsible for the short-sleeping phenotype of *appb*^-/-^ mutants.

One candidate App cleavage product that we tested was P3, as production of this peptide requires gamma-secretase but is boosted by inhibition of BACE-1. Injection of P3 also reduced nighttime sleep by affecting sleep maintenance (Figure 6). While this result underscores the potential of App-cleavage products to act as acute signals that regulate sleep, this contradicts the straightforward hypothesis that loss of P3 signaling is the cause of shortened nighttime sleep in *appb*^-/-^ mutants. However, since injection experiments do not recapitulate the precise timing or localization of P3 release, its local concentration, or (possibly) its structure, a role for P3, alone or in complex combinations with other App products like Aβ or AICD, in the observed *appb* mutant phenotypes cannot be completely ruled out. Future studies could examine whether mutations in components of γ-secretase, such as Presenilin-1 or Presenilin-2, or β-secretase, such as BACE-1, also have reduced sleep bout lengths at night. Zebrafish *bace1*^-/-^ mutants have been reported to have hypomyelination in the peripheral nervous system^25^, but to date no sleep phenotypes have been described. As we performed here for *appb* and DAPT/ lanabecestat, the examination of sleep phenotypes in *appb*^-/-^; *bace1*^-/-^ or *presenilin1/2*^-/-^ double mutants could be used to tease out which phenotypes are due to the specific cleavage of Appb.

Another possibility by which Appb could contribute to sleep regulation is raised by the recent observation that in zebrafish both Appa and Appb are colocalized to cilia and to cells lining the ventricles at 30 hours post fertilization^57^. The *appa^-/-^; appb^-/-^* double mutants were reported to have morphologically abnormal ependymal cilia and smaller brain ventricles^57^. It would be interesting to see if the localization of App proteins to cilia and ventricles are important for sleep and locomotion, as the coordinated periodic beating of the cilia is involved in the generation of CSF flow within ventricle cavities^60^, and CSF circulation is believed to facilitate transfer of signaling molecules and removal of metabolic waste products important for behaviour^61,62^.

What emerges from these results is a complex picture of endogenous App-derived signals that can regulate sleep and wake in a bidirectional manner, with some signals boosting sleep and some inhibiting sleep. Changes in the relative composition of App-derived molecules, including both Aβ42 and P3 peptides (Aβ_17-40_ and Aβ_17-42_) over the progression of preclinical and clinical AD makes for even more complex scenarios. For example, P3 peptides (Aβ_17-40_ and Aβ_17-42_) are typically found in the diffuse plaques of individuals with AD^63^, and cell culture experiments suggest that these peptides may be produced in even greater quantities than Aβ42 and Aβ40^64^. Additionally, microglia in AD patients harbor various N-terminally truncated Aβ species^65^, and the presence of P3 peptides in cerebrospinal fluid (CSF) shows a positive correlation with cognitive decline in AD patients^66^, indicating a potential role of these peptides in AD pathogenesis. Our results suggest that in addition to Aβ^26,67^, P3 might also interfere with sleep and wakefulness in AD and should be further investigated in rodent AD models.

Indicators of inadequate sleep quality, such as reduced sleep efficiency, extended time to fall asleep, heightened wakefulness during the night, and greater instances of daytime napping, have consistently been linked to compromised cognitive function ^68–71^. Furthermore, generation and release of Αβ42 into the interstitial fluid (ISF) is controlled by synaptic activity^72^ and even one night of sleep disruption can increase Aβ42 levels^73^. As sleep can directly alter Αβ levels, sleep history over one’s lifetime may be a significant contributor to AD risk and progression. Our results are consistent with the idea that alterations in App gene products may be a direct contributor to sleep phenotypes associated with preclinical and clinical AD. The specificity of the effect of Appb loss on sleep architecture also suggests that specific changes in sleep patterns that could serve as a useful AD biomarker may yet be discovered.

### Limitations of the study

We have interpreted the lack of sleep effect of the γ-secretase (or β-secretase) inhibitors in the *appb^-/-^* mutant background to be due to the loss of Appb proteolytic cleavage; however, it is technically possible that loss of Appb effects expression or localization of other γ-secretase targets (such as Notch) and the lack of drug induced sleep alterations is due to this indirect effect. Another limitation of our study is that the exogenous ventricle P3 injections might not recapitulate the actual localization or structure of P3 and further experiments would be needed to dissect the roles of the App fragments.

## METHODS

### Zebrafish strains and husbandry

Zebrafish (*Danio rerio*) were raised under standard conditions at 28°C in a 14hr:10hr light:dark cycle. All zebrafish experiments and husbandry followed standard protocols of the UCL Fish Facility. *AB*, *TL and ABxTL* wild-type strains, *appa^Δ^*^5^ *(u539)* and *appb^Δ14+^*^4^(u537) (also called *appb^-/-^* in this manuscript) were used in this study. Ethical approval for zebrafish experiments was obtained from the Home Office UK under the Animal Scientific Procedures Act 1986 under project licenses 70/7612, PA8D4D0E5, and PP6325955 awarded to JR and a personal licence 70/24631 to GGO.

#### Generation of *appa* mutant

The *appa* gene was targeted by CRISPR/Cas9 mutagenesis. CHOPCHOP (http://chopchop.cbu.uib.no) was used to identify an sgRNA to exon 18 of *appa (*ENSDARG00000104279)^31^. Cas9 mRNA was made from pT3TS-nCas9n (Addgene, plasmid 46757)^74^ using mMESSAGE mMACHINE transcription kit (Thermofisher Scientific). Constant oligomer (5’AAAAGCACCGACTCGGTGCCACTTTTTCAAGTTGATAACGGACTAGCCTTATTTTAACTTGCTATTTCT AGCTCTAAAAC-3’) and the *appa* gene-specific oligomer targeting the conserved 25-42 amino acid region of Aβ in *appa* (target sequence: 5’- GAGGACGTGAGCTCCAATAA-3’) were annealed on a PCR machine (using the program 95°C, 5 min; 95°C −>85 °C, −2°C/second; 85°C −>25°C, −0.1°C/second, 4°C) and filled in using T4 DNA polymerase (NEB) using manufacturers’ instructions at 12°C for 20 min^75^. The template was cleaned up using a PCR clean-up column (Qiaquick) and the 120 bp product was verified on a 2% agarose gel. The sgRNA was transcribed from this DNA template using Ambion MEGAscript SP6 kit^75^. 1 nl of a 1 μl of Cas9 mRNA (200 ng/μl) and 1 μl purified sgRNA (25 ng/μl) containing mixture were co-injected into one-cell stage embryos. Injected F0 embryos were raised to adulthood, fin-clipped and deep-sequenced by Illumina Sequencing (below).

#### Generation of *appb* null mutant

The *appb* gene (ENSDARG00000055543) was disrupted using TALEN mutagenesis^76^. Two TALEN arms targeting a conserved region within the first exon of zebrafish *appb* gene were designed using the Zifit software (http://zifit.partners.org/ZiFiT/)(Figure 1C)^32^ with Left Talen binding site sequence: 5’-TATGGACCGCACGGTATT-3’, Right TALEN binding site sequence: 5’-CGACTTTGTCCCTCGCCA-3’ and Spacer sequence: 5’-TTAATGCTGACGA-3’.

TALENs were generated using the FLASH assembly method following the protocol of 77. Starting with a library consisting of 376 plasmids that encode one to four TAL effector repeats consisting of all possible combinations of the NI, NN, HD or NG repeat variable di-residues (RVDs), the four 130 bp α-unit DNA fragments were amplified from each α-unit plasmid using the Herculase II Fusion DNA polymerase (Agilent) and oJS2581 and oJS2582 primers^77^. The resulting 5’ biotinylated PCR products were digested with BsaI-HF (NEB) to generate four base-pair overhangs. To generate the DNA fragments encoding the βγδε (extension fragment) and βγδ (termination fragment) repeats, each of these plasmids was digested with BbsI followed by serial restriction digests of XbaI, BamHI-HF and SalI-HF (New England Biolabs) to cleave the plasmid backbone. The four TALEN expression vectors encoding one of four possible RVDs were linearised with BsmBI (NEB). The biotinylated α unit fragments were ligated to the first βγδε fragments using Quick T4 DNA ligase and bound to Dynabeads MyOne C1 streptavidin-coated magnetic beads (Life Technologies). The bead bound α-βγδε fragments were digested with BsaI-HF (NEB) to prepare the 3’ end of the DNA fragments for the subsequent ligation step.

Each extension and termination fragment was then ligated to assemble the complete DNA fragment encoding the TALE repeat array by repeated digestion and ligation steps, and a final digestion with BbsI (NEB) released the full length fragments. The purified DNA fragments were ligated into one of four BsmBI (NEB) digested TALEN expression vectors encoding one of four possible RVDs using Quick T4 DNA ligase. Ligation products were transformed into chemically competent XL-10 Gold *E. coli* cells and clones grown on LB Agar plates containing Ampicillin at 37°C overnight. Bacterial colonies of each TALEN arm were selected and screened by colony PCR using primers oSQT34 (5′-GACGGTGGCTGTCAAATACCAAGATATG-3′) and oSQT35 (5′-TCTCCTCCAGTTCACTTTTGACTAGTTGGG-3′). Clones showing a correct sized band were cultured in LB medium containing Ampicillin at 37°C overnight. Following plasmid mini-preparation the inserts were sequenced using primers oSQT1 (5′-AGTAACAGCGGTAGAGGCAG-3′), oSQT3 (5′- ATTGGGCTACGATGGACTCC-3′) and oJS2980 (5′-TTAATTCAATATATTCATGAGGCAC-3′. mRNA was synthesised using the mMESSAGE mMACHINE T7 and polyA tailing kit. 100 pg of each of the TALEN mRNAs are injected into the cytoplasm of one-cell stage embryos, which were raised to adulthood and sequenced (below).

#### Sequencing/Genotyping Pipeline

F0 embryos were raised to adulthood, fin-clipped and deep-sequenced by Illumina Sequencing (MiSeq Reagent Nano Kit v2 (300 Cycles) (MS-103–1001)) to identify founders. Fin-clipping was done by anesthetizing the fish by immersion in 0.02% MS-222 (Tricaine) at neutral pH (final concentration 168ug/ml MS-222). DNA was extracted by HotSHOT^78^ by lysing a small piece of the fin in 50 μl of base solution (25 mM KOH, 0.2 mM EDTA in water), incubated at 95°C for 30 min, then cooled to room temperature before 50 μl of neutralisation solution (40 mM Tris-HCL in water) was added. For *appa*, a 214 base pair fragment surrounding the conserved 25-35^th^ amino acid region within *appa* was PCR amplified using gene-specific primers with miSeq adaptors (forward primer, 5’- TCGTCGGCAGCGTCAGATGTGTATAAGAGACAGCCTGCAGGAATAAAGCTGATCT-3’; reverse primer, 5’-GTCTCGTGGGCTCGGAGATGTGTATAAGAGACAG ATGGACGTGTACTGCTTCTTCC-3’). The PCR program was: 95°C – 5 min, 40 cycles of [95°C – 30 s, 60°C – 30 s, 72°C – 30 s], 72°C – 10 min. For *appb,* founders were identified by PCR amplification using the primers (forward: 5’ - TCGTCGGCAGCGTCAGATGTGTATAAGAGACAGCAG- CTGACTTTCCCTGGAGCA-3’; reverse: 5’-GTCTCGTGGGCTCGGAGATGTGTATAAGAGACAG-TGGAGGAGAACCAAGCTCCTTC-3’). PCR program was 95°C – 5 min, 40 cycles of [95°C – 30 s, 60°C – 30 s, 72°C – 30 s], 72°C – 10 min.

The PCR product’s concentration was quantified with Qubit (dsDNA High Sensitivity Assay), then excess primers and dNTPs were removed by ExoSAP-IT (ThermoFisher) following the manufacturer’s instructions. The samples were then sequenced by Illumina MiSeq to assess the presence of insertion/deletions. The mutant F0 fish containing a 5 base pair deletion in the Αβ region resulting in a stop codon on the 27^th^ residue of Αβ was chosen to generate a stable mutant line *appa^Δ^*^5^ *(u539)*. An *appb* mutant carrier containing a 14 bp deletion and a 4 bp insertion *appb^Δ14+^*^4^ (u537) that is predicted to generate a frameshift and early stop codon was selected to make stable mutant lines for further analysis. F0 fish with indels were then outcrossed to wild-types and 10 one day old F1 embryos from each pairing were screened by Sanger sequencing to assess the nature of the mutations that passed into the germline. To minimize potential off-target mutations, mutant fish were crossed to ABxTL and TL WT strains for 3 generations before performing any behaviour experiments.

#### DNA extraction

Zebrafish DNA was extracted by the HotSHOT method^78^. 50 μl of 1× base solution (25 mM KOH, 0.2 mM EDTA in water) was added to finclips in individual wells. Plates were sealed and incubated at 95°C for 30 min, cooled to room temperature and neutralised by adding 50 μl of 1× neutralisation solution (40 mM Tris-HCL in water). Genomic DNA was then stored at 4°C.

### KASP genotyping

For rapid genotyping of mutant zebrafish harbouring the *appa^Δ^*^5^ and *appb^Δ14+^*^4^ alleles, a mutant allele-specific forward primer, a wild-type allele-specific forward primer and a common reverse primer were used for KASP genotyping (LGC Genomics, KBS-1050-102, KASP-TF V4.0 2X Master Mix 96/384, Standard ROX (25mL)). The primer sequences were as follows:

*appa^Δ5^*

5’-CTTTCTCTTTGTCTCCTGCCTTCAGGTTTTCTTTGCGGAGGACGTGAGC**[TCCAA/-]**TAAAGGAGCTATTATTGGCCTGATGGTCGGAGGCGTCGTCATAGCAACCATCATCGTCATCACGCTGGTGATGCTGAGGAAGAAGCAGTACACGTCCATCCACCACGGCATCATCGAGGTGCGTGAGTTCACACCGTCTCCAC-3’

*appb^Δ14+4^*

5’-AAAATCGCGACAGAAAAACCCTGATCCGCTCAGGATATATATDCACCAGGACGT GCTGCGCTTGGGAACACAGCCATGGGTATG

GACCGCACGGTATTCCTGCT**[GTTAATGCTGACGA/TTGT]**CTTTGTCCCTCGCC ATCGAGGTAAGAATGATTGTGTAATGGAGAA

GGAGCTTGGTTCTCCTCCATACTTTAAAGGGCGGCCA-3’

where [x/-] indicates the indel difference in [WT/mutant]. 8 μl of KASP reaction is run (3.89 μl 2x KASP reaction mix (http://www.lgcgroup.com/products/kasp-genotypingchemistry/reagents/mastermix/#.VgFRWd9VhBc, 0.11 μl KBD Specific Assay (primers), 1μl H2O, 3 μl (1:10 diluted) DNA) using the protocol (94°C for 15 minutes (Activation), 94°C 20 seconds, followed by 10 cycles of touchdown PCR (annealing 61°C to 53°C, decreasing 0.8°C per cycle), 94°C 20 seconds, then 26 cycles of standard 2-step PCR at 53°C 60 seconds.

Each genotyping plate contained three wells of negative controls (KASP mastermix only with no DNA) and three wells of positive controls (DNA of a known genotype: WT, Het, Homozygous). A minimum of 24 samples were used for good clustering.

Fluorescence was read on a CFX96 Touch Real-Time PCR Detection System (Bio-Rad) and the allelic discrimination plot generated using Bio-Rad CFX Manager Software.

### Behavioural experiments

Behavioural tracking of larval zebrafish was performed as previously described^20,79^. Zebrafish larvae were raised on a 14hr:10hr light:dark cycle at 28.5°C and at were placed into individual wells of a square-well 96-well plate (Whatman) containing 650 μL of standard embryo water (0.3 g/L Instant Ocean, 1 mg/L methylene blue, pH 7.0) at 4-5 dpf. Locomotor activity was monitored using an automated video tracking system (Zebrabox, Viewpoint LifeSciences) in a temperature-regulated room (26.5°C) and exposed to a 14hr:10hr white light:dark schedule with constant infrared illumination (Viewpoint Life Sciences). Larval movement was recorded using the Videotrack quantization mode. The movement of each larva was measured, and duration of movement was recorded with an integration time of 60 sec. Data were processed using custom PERL and MATLAB (The Mathworks, R2019a) scripts, and statistical tests were performed using MATLAB (The Mathworks, R2019a).

Any one-minute period of inactivity was defined as one minute of sleep^79^. Sleep bout length describes the duration of consecutive, uninterrupted minutes of sleep whereas sleep bout number is the number of such sleep events in a given time interval. Average waking activity represents activity only during active periods.

All mutant larval zebrafish experiments were performed on siblings from *appa*^+/^*^Δ^*^5^ or *appb*^+/^*^Δ14+^*^4^ heterozygous incrosses, except for drug experiments, which were simultaneously performed on larvae from WT and *appb^Δ14+^*^4^*^/Δ14+4^* incrosses from different parents. DAPT (Cell Guidance Systems, SM15-10) was dissolved in DMSO to make a stock concentration of 10 mM and diluted further to a working concentration of 10 μM in water in 1:10 serial dilutions. 6 μl of the 10 μM DAPT stock or 6 μl of 0.1% DMSO control was added individually to the wells in the behaviour plate to make a 100 nM final concentration the second day at Zeitgeber time 0 (Lights ON) of the video-tracking in each experiment. Lanabecestat (Cambridge Bioscience, HY-100740-2mg) was dissolved in DMSO to make a to make a stock concentration of 10 mM and diluted further to a working concentration of 100 μM in water in 1:10 serial dilutions. 1.8 μl of the 100 μM Lanabecestat stock or 1.8 μl of 1% DMSO control was added individually to the wells in the behaviour plate to make a 300 nM final concentration. The drug was added on the second day at Zeitgeber time 0 (Lights ON), when the larvae are 6 dpf.

### Zebrafish *in situ* hybridization (ISH)

RNA was extracted from 30 WT embryos (5dpf) by snap freezing in liquid nitrogen and TRIzol RNA extraction (Ambion 15596026). 1 μg of RNA was reverse transcribed with AffinityScript (Agilent, 600559) to make cDNA following the manufacturer’s protocol.

Templates for *in vitro* transcription for *appa* and *appb* were generated by PCR using a reverse primer that contains a T7 promoter sequence (*appa* forward primer: 5’-CGCGGGTAAAGAGTCTGAGAGC-3’, *appa*-T7_reverse primer: 5’-<u>TAATACGACTCACTATAGGG</u> CAGACA GTATTCCTCCGACTC-3’, APPb_F: 5’-GCTCCAGGAGATATAAACGAAC-3’, APPb_T7R: 5’-<u>TAATACGACTCACTATAGGG</u> GCCGAACCTTTGGAATCTCGG-3’ using the 5 dpf cDNA library. 5 dpf larvae and 1,2,4,8 cell stage embryos were fixed in 4% paraformaldehyde (PFA) (with 4% sucrose for 5 dpf larvae) overnight at 4°C, dehydrated in a graded Methanol/PBST series (25%, 50%, 75%, 100%), kept overnight at −20°C and transferred into Phosphate Buffered Saline (PBS) the next morning. For 5dpf larvae, the brains were dissected by removing skin, cartilage, and eyes. A dioxigenin (DIG)-11-UTP-labeled antisense riboprobe targeting the gene transcript of interest was synthesized using the DIG labelling kit (Roche) and T7 RNA polymerase^80^. Probe hybridization was carried out in hybridization buffer (50% formamide (v/v), 5x SSC (750 mM NaCl, 75 mM sodium citrate), 9.2 mM citric acid, 0.5 mg/mL Torula RNA, 0.05 mg/mL heparin, 0.1% Tween-20) supplemented with 5% dextran sulfate overnight at 65°C. The larvae were then incubated overnight at 4°C with anti-Dig-AP (1:2000 in 5% normal goat serum, 11093274910, Roche) and washed before detecting alkaline phosphatase using NBP/BCIP(Roche) according to the manufacture’s guidelines.

### *in situ* Hybridisation Chain Reaction (HCR)

#### Probe design

*In situ* Hybridization chain reaction (HCR) probes were designed as described^81^. HCR split initiator sequences B1, B3 and B5 were taken directly from Choi et al. (2018)^81^. To generate optimal probe sets, probe pairs were excluded if they fell below the pre-set melting temperature and % GC thresholds. Probe pairs with strong sequence similarity to off-target transcripts were also excluded. For both *appa* and *appb* transcripts we generated probe sets that contained 20 probe pairs. HCR multiplexing was used to stain both paralogs in the same brain. HCR probes sets were purchased as custom DNA oligos from Thermo Fisher Scientific (UK), and HCR Amplifiers (B1-Alexa Fluor 488, B3-Alexa Fluor 546, and B5-Alexa Fluor 647) and buffers were purchased from Molecular Instruments (Los Angeles, CA, USA).

##### Probe info

| Gene | Transcript ID | Initiator | # probe pairs per set |
| --- | --- | --- | --- |
| *appa* | ENSDART00000170483.2 | B3 | 20 |
| *appb* | ENSDART00000077901.5 | B5 | 20 |

### *In situ* hybridization chain reaction

In situ hybridization chain reaction (HCR) was performed following the “HCR RNA-FISH protocol for whole-mount zebrafish embryos and larvae” from Choi et al., 2018^81^. The protocol was modified by omitting all proteinase K and methanol permeabilization steps. HCR multiplexing was performed by pooling probe sets targeted to *appa, appb*, and *gad1b* (*gad1b* was necessary to register brains to the zbb atlas, http://zbbrowser.com). WT 6dpf larvae that had been raised in 10hr:14hr normal or reversed light: dark cycle were euthanised post 4 hours lights on (for day condition) or lights off (for night condition) and fixed in 4% paraformaldehyde with 4% (w/v%) sucrose overnight at 4°C. Larvae were removed from the fixative by washing three times for 5 min in PBS before being transferred to a SYLGARD coated dish, where the eyes and the skin that covers the brain were removed with a pair of sharp forceps. After dissection, a brief 20 minute postfix in 4% paraformaldehyde was performed followed by 3 further washes in PBST (1× DPBS + 0.1% Tween 20) to remove any fixative. Sample preparation was completed by two brief PBST washes. For the probe hybridization stage, we first incubated larvae in hybridization buffer in a heat block at 37°C for 30 min. The *appa* and *appb* probe solution was prepared by adding 4 µl of each 1 µM probe set to 500 µl hybridisation buffer (final probe concentration of 4 pmol). Hybridization buffer was replaced by the probe solution, and larvae were incubated at 37°C overnight (12–16 hr).

The next day, excess probe were removed by washing larvae four times 15 min in probe wash buffer preheated to 37°C, followed by two 5 min washes in 5x SSCT (5× sodium chloride sodium citrate + 0.1% Tween 20) at room temperature. Samples were kept at room temperature for subsequent amplification steps. Larvae were transferred to room temperature amplification buffer for 30 min. 3 µM stocks of hairpin H1 and hairpin H2 were individually snap cooled by heating to 95°C for 90 seconds then left to cool to room temperature in a dark drawer for 30 min. A hairpin solution was prepared by adding 10 µL of hairpin H1 and 10 µl of hairpin H2 to 500ul amplification buffer (final hairpin concentration of 30 pmol). Finally, pre-amplification buffer was removed and the hairpin solution was added to the larvae that were incubated overnight (12–16 hr) in the dark at room temperature. After overnight incubation, excess hairpins were removed by washing in 5× SSCT for 2 x 5 minute, then 2 x 30 minutes and finally 1 x 5 minute at room temperature. Larvae were transferred to PBS and kept at 4°C protected from light for up to 3 days.

### Imaging

Following the HCR staining protocol, larvae were imaged using a ZEISS Lightsheet Z.1, fitted with a ZEISS W Plan-APOCHROMAT 10x/0.5 detection objective and two ZEISS LSFM 5x/0.1 illumination objectives. Larvae were mounted in glass capillary tubes filled with 1% low-melting agarose and held vertically, anterior facing up in a ZEISS Lightsheet Z.1 sample holder. HCR Amplifiers tagged with Alexa Fluor 488 (*gad1b*), Alexa Fluor 546 (*appa*), and Alexa Fluor 647 (*appb*) were imaged using 488 nm, 561 nm and 638 nm lasers, respectively. Full brain stacks were acquired with a final image size of 1920 x 1920 pixels (889.56 um x 889.56 um) and a voxel size of 0.46 um x 0.46 um x 1.00 um.

### Registration

To allow for comparison of *appa* and *appb* expression among fish, all brains were registered onto the zbb brain atlas (http://zbbrowser.com) using the ANTs toolbox version 2.1.0^82^. Raw Zeiss CZI files were rescaled, converted to 8-bit grayscale and saved in the NRRD file format. The *gad1b* channel from each *gad1b*-*appa*-*appb* hyperstack was registered to a *gad1b*-HCR reference brain (ref_gad1b) (1 × 1 × 1 xyz µm/px) using the following ANTs script:

fileext= nrrd

float= F1_app (e.g.)

ref= ref_gad1b (e.g.)

addstr = zbb (e.g.)

interp=BSpline

rigidm=GC

affinem=GC

synm=CC c

synp=[0.05,6,0.5]

antsRegistration -d 3 --float 1 -o [${float}_01${addstr}_,${float}_01${addstr}_Warped.nii.gz] -n ${interp} -r [${ref},${float}_01.${fileext}, 0] -t Rigid[0.25] -m ${rigidm}[${ref},${float}_01.${fileext},1,32,Regular,0.25] -c [200×200×200×0,1e-8,10] -f 12×8×4×2 -s 4×3×2×1 -t Affine[0.25] -m

${affinem}[${ref},${float}_01.${fileext},1,32,Regular,0.25] -c [200×200×200×0,1e-8,10] -f 12×8×4×2 -s 4×3×2×1 -t SyN${synp} -m ${synm}[${ref},${float}_01.${fileext},1,2] -c [200×200×200×200,1e-7,10] -f 12×8×4×2 -s 4×3×2×1

The transformation files were applied to the *appa* and *appb* channels using the

following ANTs script:

antsApplyTransforms -d 3 -v 0 --float -n ${interp} -i ${float}_0${i}.${fileext} -r ${ref} -o

${float}_0${i}${addstr}_Warped.nii.gz -t ${float}_01${addstr}_1Warp.nii.gz -t

${float}_01${addstr}_0GenericAffine.mat

#### Western Blots

Protein was extracted from adult *appa ^Δ^*^5^*^/ Δ^*^5^; *appb*^-/-^ mutants or WT larvae. Fish were euthanized, brains were dissected out and immediately frozen in liquid nitrogen prior to use and stored at − 80°C. Samples were homogenized in an ice-cold lysis buffer (10 mM Tris–HCl pH 8.0, 2% sodium deoxycholate, 2% SDS, 1 mM EDTA, 0.5 M NaCl, 15% glycerol) supplemented with protease inhibitors cocktail (Calbiochem protease inhibitors cocktail III) using a syringe needle (BD microlance, Ireland; 27G ½” 0.4 x 13 mm) on ice. Samples were then incubated 20 min on ice, sonicated for 10 min at 70% amplitude with a pulse of 30s on and off and then centrifuged at 10,000×*g* at 4°C. Supernatants were collected and kept on ice and protein concentration measured with a Qubit™ protein BR assay kit (Thermo Fisher Scientific, Waltham, MA) and samples stored at − 80°C. Protein samples (40-50ug) were then diluted in a denaturing lysis buffer (1X NuPAGE LDS Sample Buffer (Thermo Fisher Scientific, Waltham, MA), 0.05 M DTT (Sigma-Aldrich, St. Louis, MO), lysis buffer completed with protease inhibitors) and then boiled for 5 min at 95°C. Proteins were then separated on a NuPAGE NOVEX 4-12% gradient Bis– TRIS pre-cast gel (Thermo Fisher Scientific, Waltham, MA) at 150V for 1 hour and transferred onto a 0.2 μm Amersham™ Portran™ nitrocellulose membrane at 400 mA for 50 minutes on ice. The membrane was incubated in a blocking solution (5% milk) for 2 hours at RT and then immunoblotted overnight at 4°C with the primary mouse anti-amyloid precursor protein A4 antibody (clone 22C11) (Sigma MAB348-AF647) (1:3000) and with a loading concentration control mouse anti-γ-tubulin monoclonal (1:10,000) (Sigma, St. Louis, MO). The membrane was then washed in TBS-Tween three times for 10 min at RT and incubated with the secondary antibody anti-mouse-HRP (1:5000) (Cell Signaling, Danvers, MA) for one hour at RT. The signal was developed using SuperSignal West Dura Extended Duration Substrate kit (Thermo Fisher Scientific, Waltham, MA) and imaged using ChemiDoc Imaging (Bio-Rad, Hercules, CA). Western blot images were processed using ImageJ (NIH, USA).

### qPCR

Total RNA was isolated from 4-6 dpf zebrafish larvae using the RNeasy Plus Micro Kit (Qiagen). cDNA was synthesized using the SuperScript III First-Strand Synthesis System (Invitrogen). qPCR was performed using a CFX96 machine (Bio-Rad) and accompanying BioRad CFX Manager (v3.1) using GoTaq qPCR master mix (Promega, A6001) with the primers qPCR_appb_F2 (5’-CGTGGTCATCGCTACTGTCA) and qPCR_appb_R2 (5’-CTGCCGCATCCACCTCAATA) at 60°C resulting in a 98 bp product. *ef1α* was chosen as the reference gene as it has been validated in the zebrafish for qPCR normalisation^83^. Reactions were performed up to a total volume of 10µl per reaction with primer concentrations of 10µM for *ef1a, (eef1a1*_qRT-PCR Forward: 5’-TGCTGTGCGTGACATGAGGCAG-3’ and *eef1a1_*qRT-PCR Reverse: 5’-CCGCAACCTTTGGAACGGTGT-3’) or 20µM for *appb* primers. Efficiency of *ef1α* and *appb* primers were established to be within acceptable efficiency thresholds through a serial dilution series (90-110% efficiency). Threshold cycle values (Cq) were obtained for each gene in each sample in technical replicates. Two replicate experiments were performed per gene with 3 technical replicates.

### qPCR analysis

Analysis was undertaken using the ‘delta-delta Cq’ method to compare the relative gene expression of the target gene (*appb*) to the reference gene (*ef1α*)^84^. Further analysis used the Qiagen REST programme (2009)(v2.0.13). This software compares treated versus untreated samples using provided serial dilution data for efficiency calculations. REST then performs a pairwise fixed reallocation randomisation test (permutations = 2000) to determine p values between samples, as well as giving confidence intervals and standard error of the permutation analysis. Melt curve analysis was conducted through the BioRad CFX manager software. Boxplots were generated using GraphPad Prism (Dotmatics).

#### Endogenous Aβ measurements in adult zebrafish brains

For the DAPT treatment, WT adult fish were treated with a final concentration 25 μM of DAPT for 24 hours in a small tank. Adult zebrafish brains were dissected, weighed and were mechanically homogenized in 100 μL TBS (50mM Tris-HCL, pH 8.0) containing Calbiochem protease inhibitor cocktail set III (1:200). Whole brain homogenates were centrifuged at 16,000 g at 4°C for 30 min, the supernatant was aliquoted and stored at −80°C. Total protein concentration of the samples was determined using a Pierce Detergent Compatible Bradford assay kit according the manufacturer’s instructions (Thermofisher, 23246). Aβ40 and Aβ42 measurements of the samples were done according to the manufacturers protocols using Mesoscale Discovery V-plex Plus Aβ42 (4G8) or V-plex Plus Aβ peptide panel 1 (4G8) kits on the Meso Scale Discovery platform (MSD, Rockville, Maryland) in technical duplicates or triplicates. Standard curves were created using the MSD Mesoscale Discovery Workbench Toolbox to benchmark Aβ concentrations in the samples. A 4-parameter logistic curve was used to fit standards and calculate the concentration for unknowns and Aβ controls. Aβ standards for the calibration curve were measured in duplicate and were set in serial in 1:4 dilutions. The upper and lower limits of detection were set as 2.5 standard deviations from the bottom and top calibrator. The calculated Aβ amounts were then normalized to the total extracted protein levels from each sample were determined using the Bradford assay.

#### P3 (Beta-Amyloid 17-42 peptide) preparation and injection

HFIP-treated Beta-Amyloid 17-42 peptide (P3) (Cambridge Biosciences) was dissolved in DMSO and vortexed briefly to yield a 100μM solution. The stock solution was aliquoted as 5μl in individual tubes and kept at −80°C. Just before the injections, stock solutions were 1:10 serially diluted in Phosphate buffered saline (PBS) to obtain a 10 nM solution. The Injections were carried out with a Pneumatic PicoPump (WPI) and glass capillary needles (Science Products Gmbh) prepared with a Micropipette Puller (Shutter Instruments). Larvae (5 dpf) were anesthetized using 4% Tricaine (42 mg/L, Sigma) 30 minutes before injections. Larvae were immobilized dorsal up in 2% low melting point agarose (Thermo Fischer) in fish water on a small petri dish lid. 1 nL of the 10 nM Beta-Amyloid 17-42 peptide solution was injected into the hindbrain ventricle. For controls, 1nL of 1x PBS was injected instead. Success of all injections was confirmed by judging the inflation of the ventricles. Larvae were removed from the agar, put into fresh water for 20 minutes to recover from the Tricaine, and transferred into a 96 square-well plate to undergo sleep/wake behaviour tracking (see **Methods**: **Behavioural experiments)**.

#### Statistical Tests

Effect sizes were calculated using the dabest estimation package implemented in MATLAB^85^ by creating multiple bootstrap *resamples* from the dataset, computing the effect sizes on each of these resamples, and determining the 95% CI from these bootstrapped resamples^85^. All other statistical tests were done using MATLAB (The Mathworks, R2019a). Data was tested first for normality using the Kolmolgorov-Smirnov Normality test, outliers were detected removed using Grubb’s test at p<0.01, and the data was analysed with one-way ANOVA or Kruskal-Wallis followed by Tukey’s or Dunnett’s post-hoc test. A two-way ANOVA was used to calculate the interaction statistics for the drug x genotype experiments.

## Acknowledgements

We thank Alexandra Abramsson and the Zetterberg Lab for providing antibodies, Chintan Trivedi for helping with probe design and MATLAB scripts for brain registration, Leonardo Valdivia and Karin Tuschl for the TALENS, UCL Fish Facility for animal husbandry, Shreena Nayee for genotyping and fish maintenance, Laura Roesler Nery, and members of the Rihel lab and the UCL Zebrafish 1^st^ floor for experimental/technical support and useful discussions.

This work was funded by Wellcome Trust Investigator Award (#217150/Z/19/Z), BBSRC Research Grant (#BB/T001844/1), and an Alzheimer’s Research UK Interdisciplinary Grant to JR and by a UCL Neuroscience Domain Grant and an Alzheimer’s Research UK Research Fellowship (ARUK-RF2022B-015) to GGO.

## Author Contributions

G.G.O conceptualized and designed the experiments, generated mutants, performed the *appb* behaviour baseline and *appb* drug-interaction experiments, endogenous amyloid beta measurement experiments, analysed the data and wrote the paper; S.L performed the *appa* baseline behaviour experiments; T.C. performed the *appb* qPCR experiments; L.T. performed the P3 injection experiments; J.D. performed the HCR experiments, T.K. performed the Western Blot experiments; J.R. conceived of and designed experiments, supervised the project, and wrote and edited the paper.

---

**Supplemental Figure S1:**
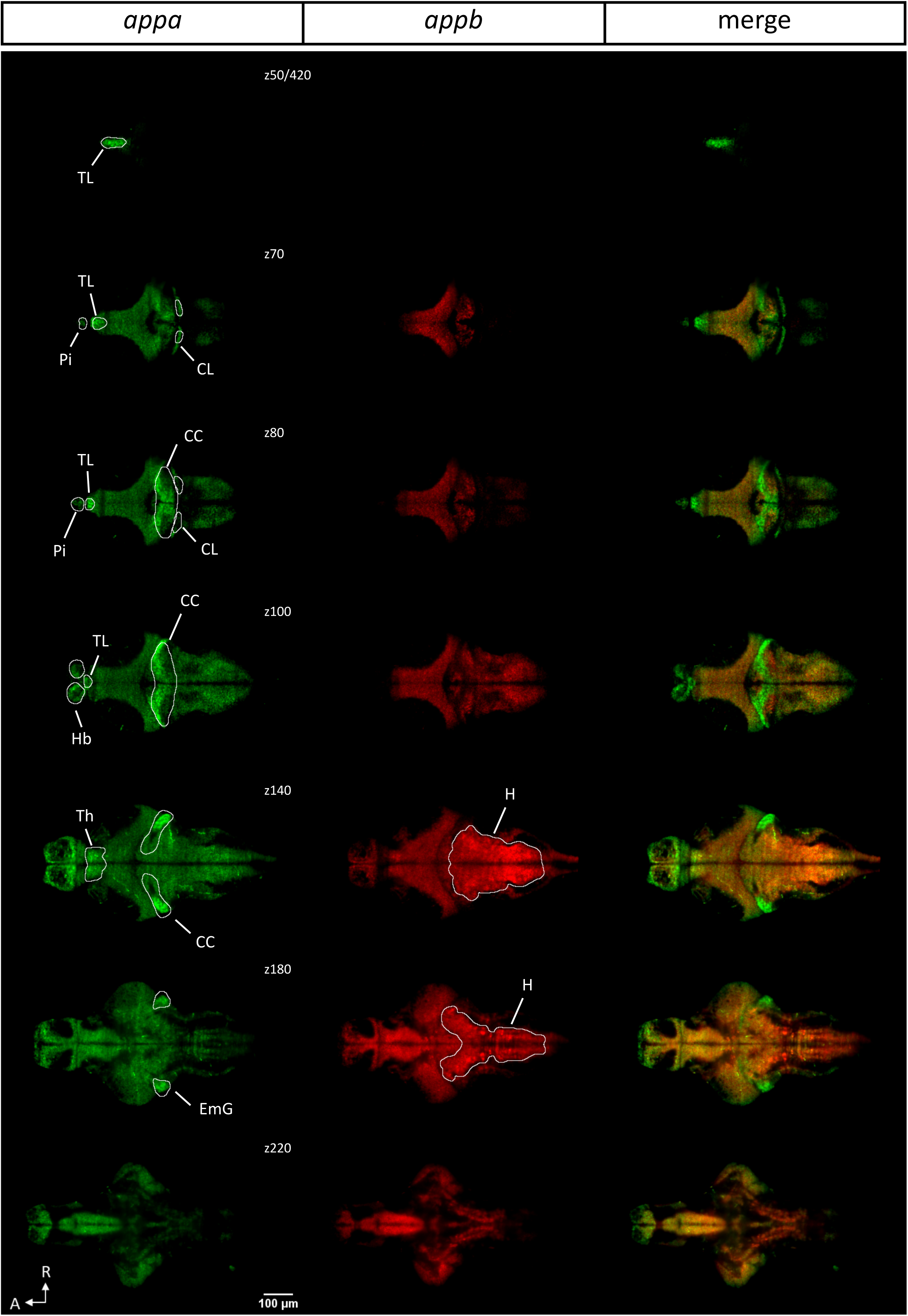
*appa* and *appb* have differential brain expression. As detected by multiplexed hybridization chain reaction (HCR), *appa* (green) and *appb* (red) are expressed differentially at 5 dpf. Dorsal view of several individual z-planes (z50, 70, 80, 100, 140, 180, 220) from a single representative brain (n=22 larvae) are shown. Brains were registered to the ZBB brain atlas using the *gad1b*-HCR reference brain, and outlines of anatomical regions are based on the Z-brain binary masks (Randlett et al., 2015). For the habenula, torus longitudinalis, and pineal gland, the outlines were translated to account for suboptimal registration. The hindbrain outline was generated manually. TL = Torus longitudinalis, Pi = Pineal, CL = Cerebellum – caudal lobe, CC = Corpus cerebelli, Hb = Habenula, EmG = Eminentia granularis, H = Hindbrain; A = Anterior, R = right.

**Supplemental Figure S2:**
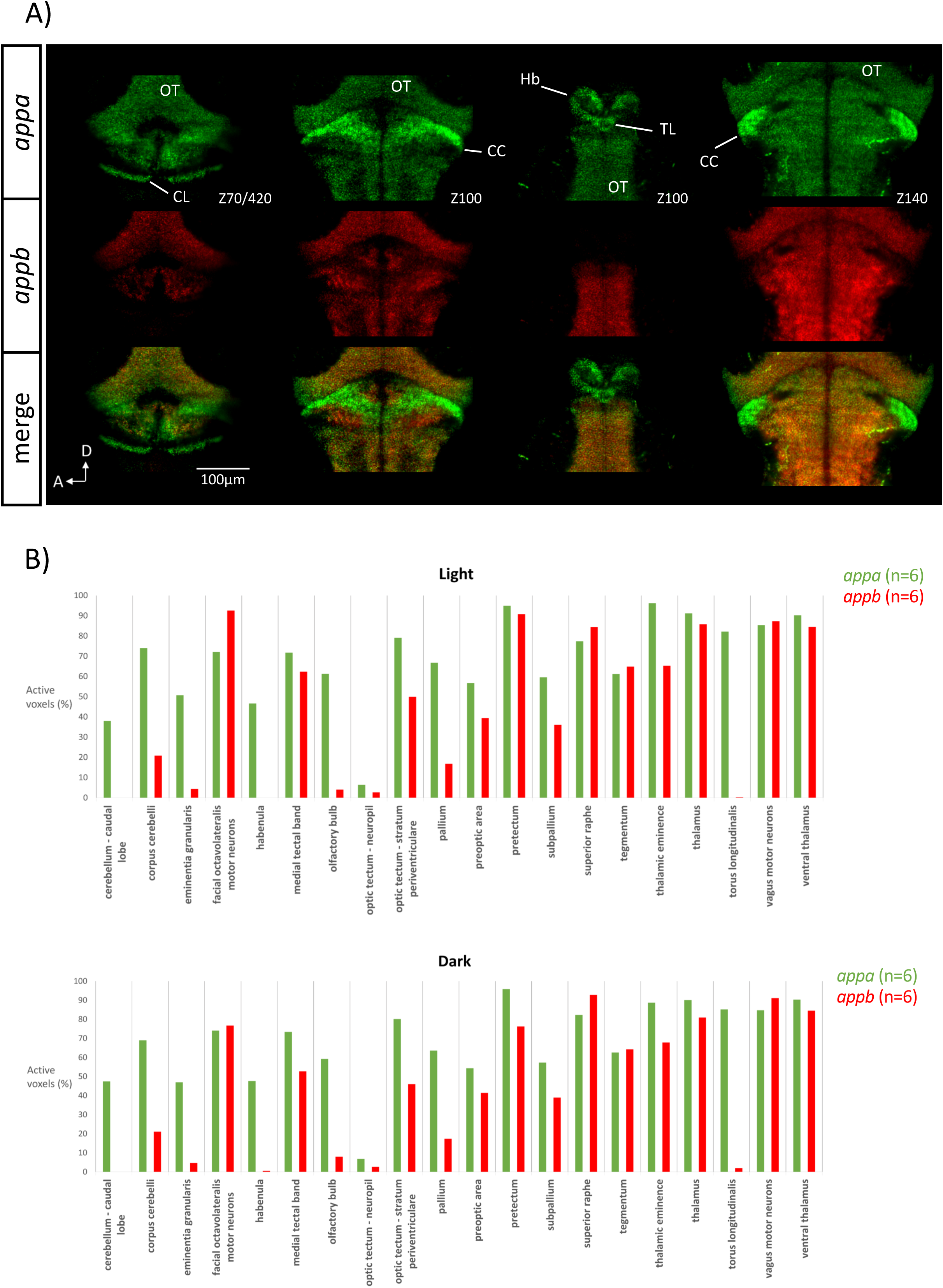
*appa* and *appb* brain expression. A) Cropped and expanded dorsal views showing *appa* (green) and *appb* (red) expression visualised with multiplexed HCR. Several individual z-planes (z70, 100, 140) from a single representative brain (n=22) are shown. Some weak expression of *appb* in the habenula is not visible due to thresholding. OT= Optic Tectum, TL = Torus longitudinalis, CL = Cerebellum – caudal lobe, CC = Corpus cerebelli, Hb = Habenula; A = anterior, D = Dorsal. B) Mean % active voxel values of multiplexed in situ hybridisation signals for *appa* (green) and *appb* (red) presented from light (top panel, n=6) & dark (bottom panel, n=6) conditions across brain regions.

**Supplemental Figure S3:**
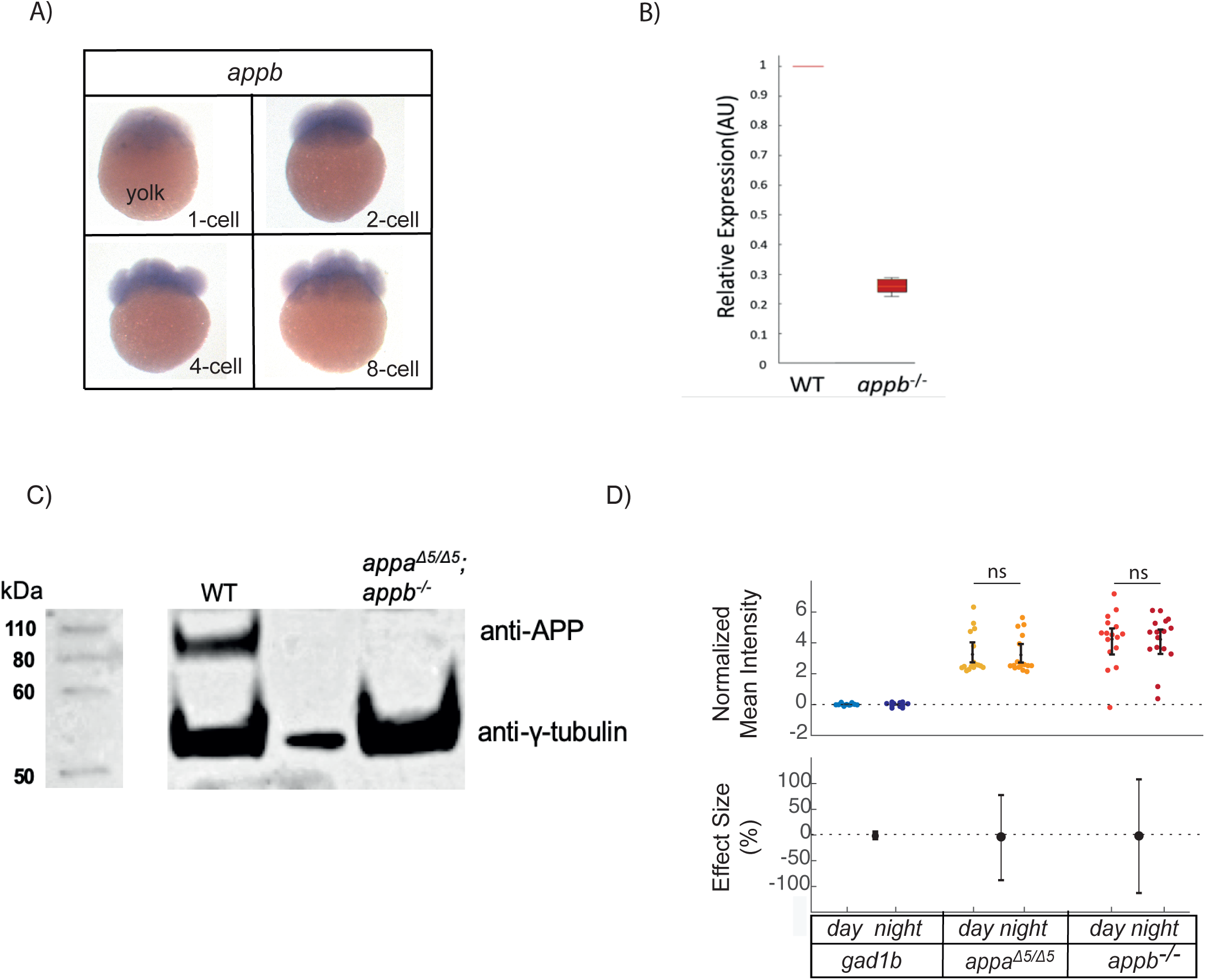
*appa* and *appb* expression. **A)** As detected by in situ hybridization (ISH), *appb* is expressed in early development (1-8 cell stage, dark purple staining), indicating that it is maternally deposited. **B)** qRT-PCR of *appb* transcript levels from 5dpf WT and *appb^-/-^* mutant larvae, indicating non-sense mediated decay. **C)** The uncropped Western blot analysis from Figure 1E of APP in brain homogenates from wildtype (WT) and *appa^Δ5/Δ5^; appb^-/-^*double mutants. An empty column was left in between samples to avoid spill-over, but was cropped in Figure 1E for clarity. **D)** Mean intensity of *appa* and *appb* expression visualized with multiplexed in situ hybridisation, normalized to the reference channel *gad1b* at 5 dpf during the day and night. Each dot represents one brain image. N=16 per condition.

**Supplemental Figure S4.**
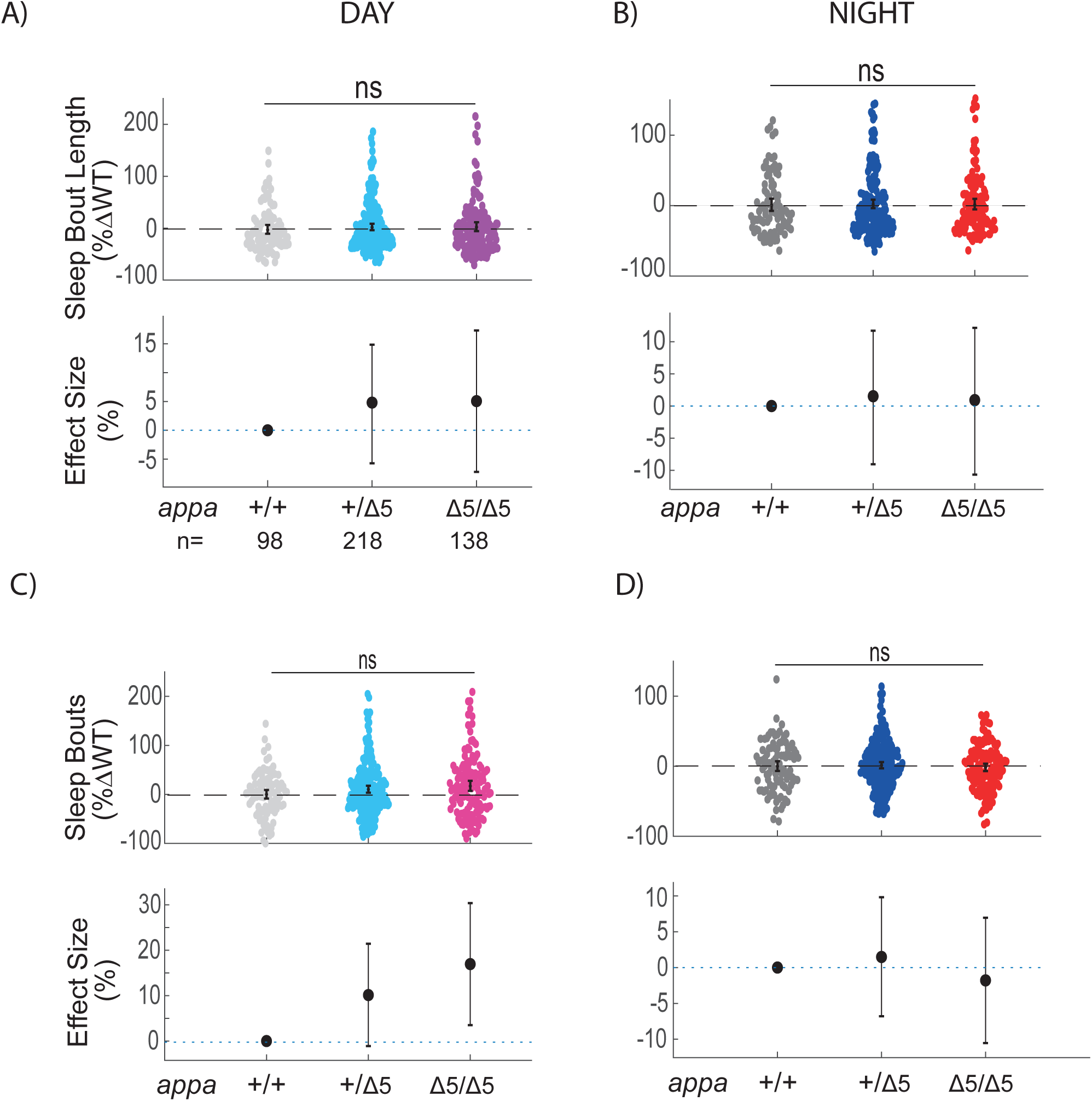
Sleep parameters of *appa* mutants. Day sleep length, **B)** Night sleep length, **C)** day sleep bout number and **D)** night sleep bout number of *appb^-/-^* mutants normalized to WT siblings across N=5 independent experiments. At top, each animal (dots) is normalized to the mean of their experimentally-matched WT data. At bottom, the effect size ± 95%CI relative to the WT mean are shown. ^ns^p>0.05, Kruskal-Wallis followed by Tukey’s test. n = the number of larvae.

**Supplemental Figure S5.**
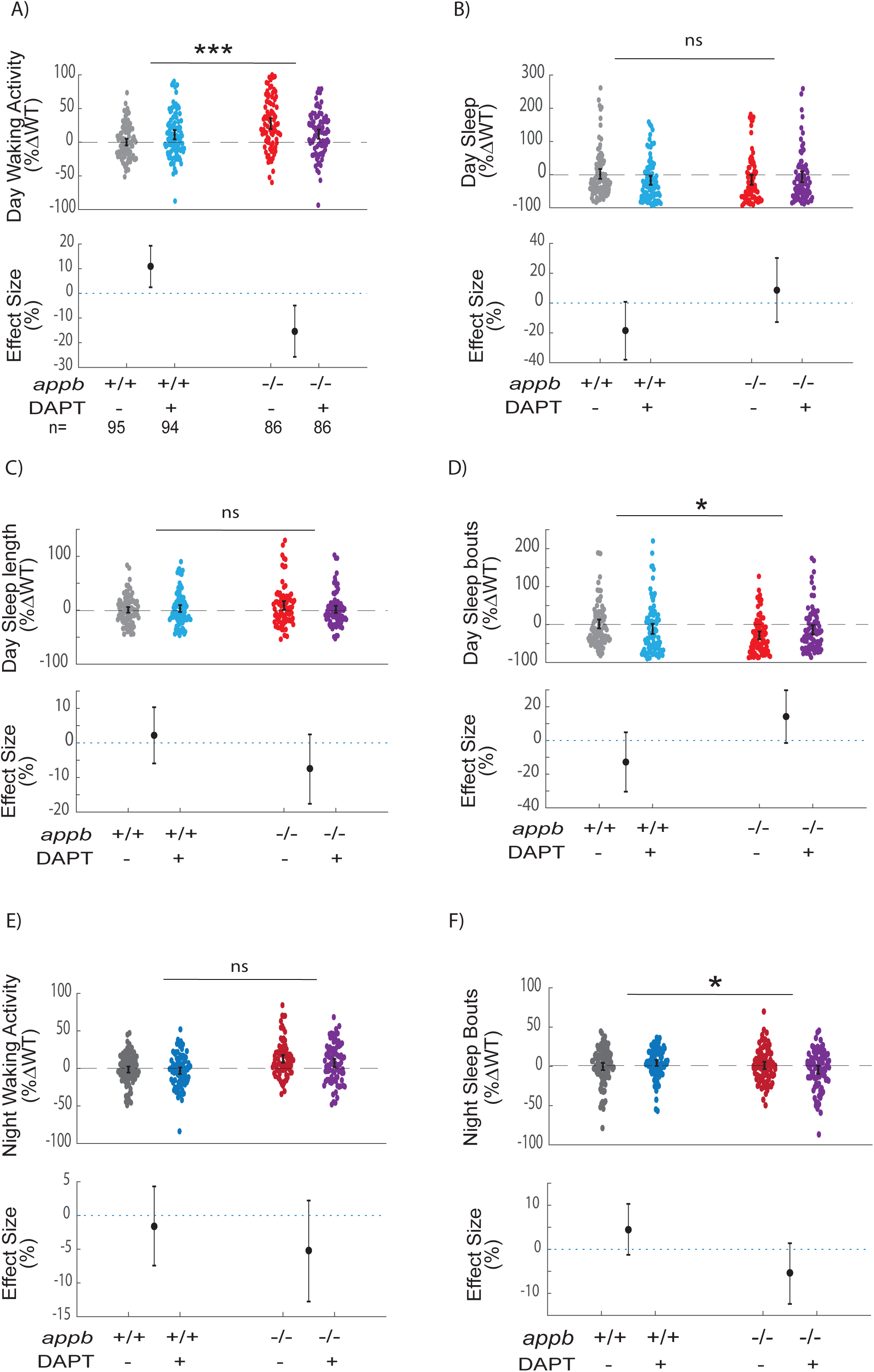
The y-secretase inhibitor DAPT affects multiple behaviors in WT and *appb*^-/-^ mutants. **A)** Average day waking activity, **B)** day sleep, **C)** day sleep bout length, **D)** day sleep bout number, **E)** night waking activity, and **F)** night sleep bout number of WT and *appb^-/-^* mutants exposed to either DAPT or DMSO vehicle. At top, each dot represents a single larva normalized to the mean of the experimentally-matched WT fish in DMSO, and error bars indicate ± SEM. At bottom, the effect size and 95%CI are plotted. n = the number of larvae. Data is pooled from N =4 independent experiments. p>0.05, *p≤0.05, **p≤0.01, 2-way ANOVA. p values indicate drug x genotype interaction.

**Supplemental Figure S6.**
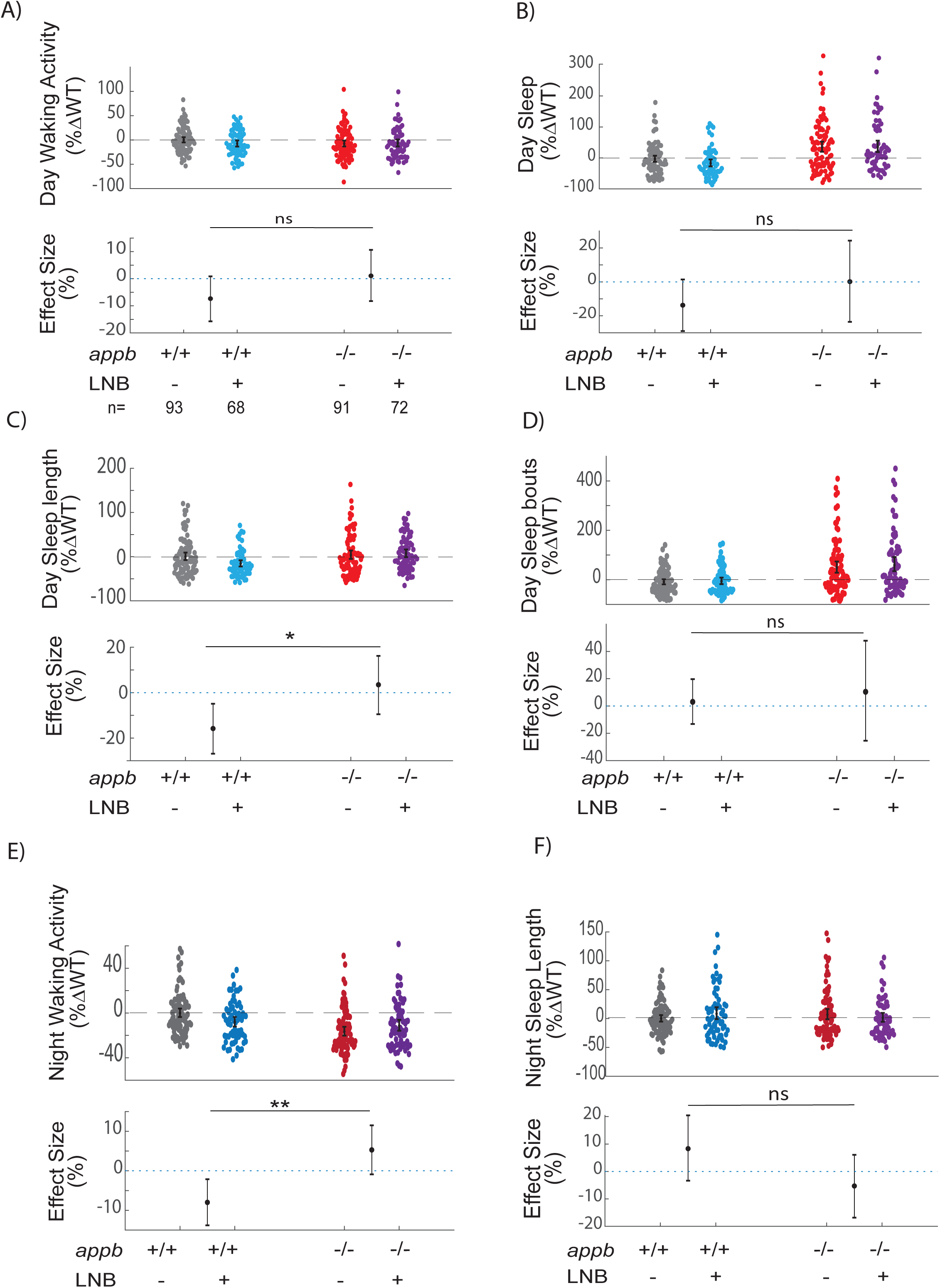
The β-secretase inhibitor Lanabecestat alters sleep at night in WT but not in *appb*^-/-^ mutants. **A)** Average day waking activity, **B)** day sleep, **C)** day sleep bout length, **D)** day sleep bout number, **E)** night waking activity, **F)** night sleep bout number of WT and *appb^-/-^* mutants exposed to 0.3 µM Lanabecestat or DMSO vehicle. At top, each dot represents a single larva normalized to the mean of the experimentally-matched WT fish in DMSO, and error bars indicate ± SEM. At bottom, the effect size and 95%CI are plotted. n = the number of larvae. Data is pooled from N =4 independent experiments. p>0.05, *p≤0.05, **p≤0.01, 2-way ANOVA. p values indicate drug x genotype interaction.

